# Oxidative Injury to Lung Mitochondrial DNA is a Key Contributor for the Development of Chemical Lung Injury

**DOI:** 10.1101/2024.11.22.624949

**Authors:** Shubham Dubey, Zhihong Yu, Emily Morgan Stephens, Ahmed Lazrak, Israr Ahmad, Saurabh Aggarwal, Shaida Andrabi, M. Iqbal Hossain, Tamas Jilling, Solana R. Fernadez, Jennifer L. Bartels, Suzanne E. Lapi, James Mobley, Viktor M. Pastukh, Mark Gillespie, Sadis Matalon

## Abstract

The mechanisms and extent to which inhalation of oxidant gases damage the mitochondrial genome contributing to the development of acute and chronic lung injury have not been investigated. C57BL/6 mice exposed to chlorine (Cl_2_) gas and returned to room air, developed progressive loss of lung DNA glycosylase OGG1, significant oxidative injury to mtDNA, decreased intact lung mitochondrial (mt) DNA, generation of inflammatory pathway by DAMPs causing airway and alveolar injury with significant mortality. Global proteomics identified over 1400 lung proteins with alteration of key mitochondrial proteins at 24 h post Cl_2_ exposure. Intranasal instillation of a recombinant protein containing mitochondrial targeted OGG1 (mitoOGG1) post exposure, decreased oxidative injury to mtDNA, lung mitochondrial proteome, severity of the acute and chronic lung injury and increased survival. These data show that injury to the mt-genome is a key contributor to the development of acute and chronic lung injury.

## Introduction

Global production of chlorine (Cl_2_) exceeds millions of tons per year. Uses of Cl_2_ include pulp bleaching, waste sanitation, chemical and pharmaceutical manufacturing, drinking and recreational water treatment. Between 1940 and 2007, the accidental release of Cl_2_ in 30 large cities worldwide (52) and the deliberate release of Cl_2_ during acts of terrorisms by insurgents in Iraq and Syria (15) caused significant mortality and morbidity to humans and animals (3, 15, 16, 26, 51). Humans exposed to 600 ppm for 30 minutes (min.) developed Acute respiratory Distress syndrome (ARDS) and required tracheal intubation and mechanical ventilation(52). Survivors are more prone to bacterial infections and the development of pulmonary fibrosis (2). There are no safe exposures to Cl_2_: Exposures to swimming pool water treatment (13) and household accidents due to mixing bleach with acidic cleaners have resulted in wheezing and exacerbate the asthma and pulmonary diseases (20). Current treatment for toxic Cl_2_ gas exposure is supportive (15) but targeted therapies are unavailable.

Similar to typical ARDS, Cl_2_ gas-induced lung injury is associated with a profound oxidative stress (5, 28, 53, 54) detectable in rodents, human subjects and human cells during and post exposure to Cl_2_ (4, 5, 28). The mitochondrial genome is critical for the maintenance of bioenergetics and mitochondrial signaling activities and is highly sensitive to oxidative stress (21, 22). The contribution of oxidative mitochondrial damage to inflammatory lung diseases is undisputed, but mechanisms underlying this association remain to be fully elucidated.

While mtDNA damage is directly cytotoxic, accumulating evidence shows that mtDNA damage-dependent DAMP formation also plays a critical pathogenic role by activating downstream inflammatory cascades that could propagate injury to remote sites even if they are unaffected by the initial insult (45, 49, 57). This may be critical to the delayed effects of Cl_2_ gas where the inflammatory pathways activated by mtDNA DAMPs engage feed-forward inflammatory pathways that could propagate injury despite control of the initiating stimuli (29, 36, 38, 41, 42).

We therefore, tested the hypothesis that an early molecular target in Cl_2_ gas inhalation could be the mt-genome, in which oxidative damage may trigger bioenergetics dysfunction as well as generation of DAMPs that drive acute and chronic lung injury. Further, we hypothesize that repair of mt-genome by administering a mitochondrial-targeted recombinant form of the DNA glycosylase OGG1 (mitoOGG1) post-exposure, may restores bioenergetics, reduces inflammation and mitigate or repair the onset and development of acute and chronic lung injury in rodents.

To accomplish these goals, we exposed adult C57BL/6 male and female mice to Cl_2_ gas in concentrations and times likely to be encountered in the vicinity of industrial accidents and returned them to room air. We first performed a series of biochemical measurements to assess oxidative injury to mitochondrial and nuclear DNA, and the extent of inflammatory responses as shown by the activation of mitochondrial DAMP receptors, such as NLRP3, TLR9, and concentrations of their downstream cytokines and chemokines in the plasma. We then assessed physiological injury to lung airways and blood gas barrier, as well as specific injury to lung alveolar ATII and ATI ion channels both ex vivo, by patching cells in lung slices in the cell attached mode and in vivo by measuring sodium-driven clearance of alveolar fluid post exposure. Selective measurements were also repeated in mice with global (mitochondrial and nuclear) OGG1 depletion (OGG1^-/-^).

To establish causative relationships among oxidative mitochondrial injury and the onset and development of acute and chronic lung injury we repeated these measurements following instillation of a fusion protein containing mitochondrial targeted OGG1 (mitoOGG1) in mice post Cl_2_-exposure, assessed its pharmacokinetics by radiolabeling with ^89^Zr, and the extent to which it repaired the mtDNA oxidative stress, acute and chronic lung injury and improved survival. Additionally, we performed unbiased high throughput proteomics mass spectrometry in lung tissues to identify novel mitochondrial specific targets that were changed with Cl_2_ and subsequently reversed/attenuated with mitoOGG1. Finally, to assess whether exposure to Cl_2_ damages key mitochondrial functions, we exposed human airway cells to Cl_2_ gas and assessed the extent to which treatment with mitoOGG1 repaired the Cl_2_ induced injury to their bioenergetics as measured by analyzing the differential mitochondrial oxygen consumption rates. These integrated physiological, biochemical, molecular cell biology and biophysical studies in wild type and OGG1^(-/-)^ mice allowed us to clearly identify the mechanisms by which oxidative injury to lung mitochondria contributes to the development of acute and chronic lung injury.

## Results

### Exposure of C57BL/6 mice to Cl_2_ (500 ppm for 30 min) downregulates lung OGG1 after they were returned to room air

Exposure of C57BL/6 mice to Cl_2_ significantly downregulates the expression of lung OGG1 at both mRNA and protein levels, starting at 6h post exposure (**Fig. 1A, B, C**). At 24 h post exposure lung OGG1 mRNA was decreased by more than 90% while OGG1 protein levels were decreased by 50%. Intact lung mtDNA (assessed by RT-PCR) decreased at 1, 6 and 24 h post Cl_2_ (**Fig. 1D,E**) with a concomitant increase of lung mtDNA lesions (**Supplementary** Fig. 1A). Southern blot analysis of lung mtDNA in the presence and absence of Fpg (**Fig. 1F**), a base excision repair enzyme which recognizes and removes oxidized purines from damaged DNA confirmed the presence of mtDNA injury.

**Figure 1.**
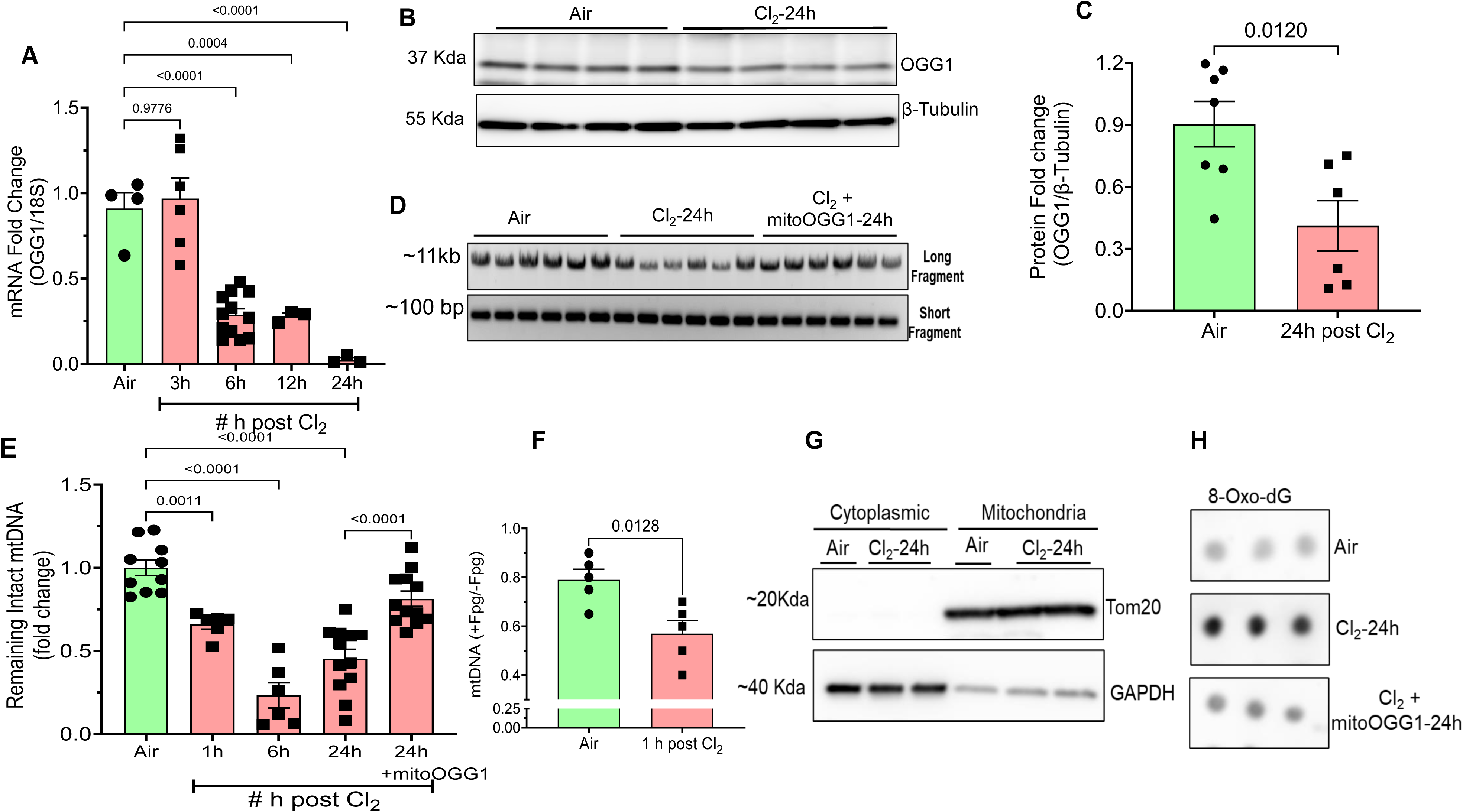
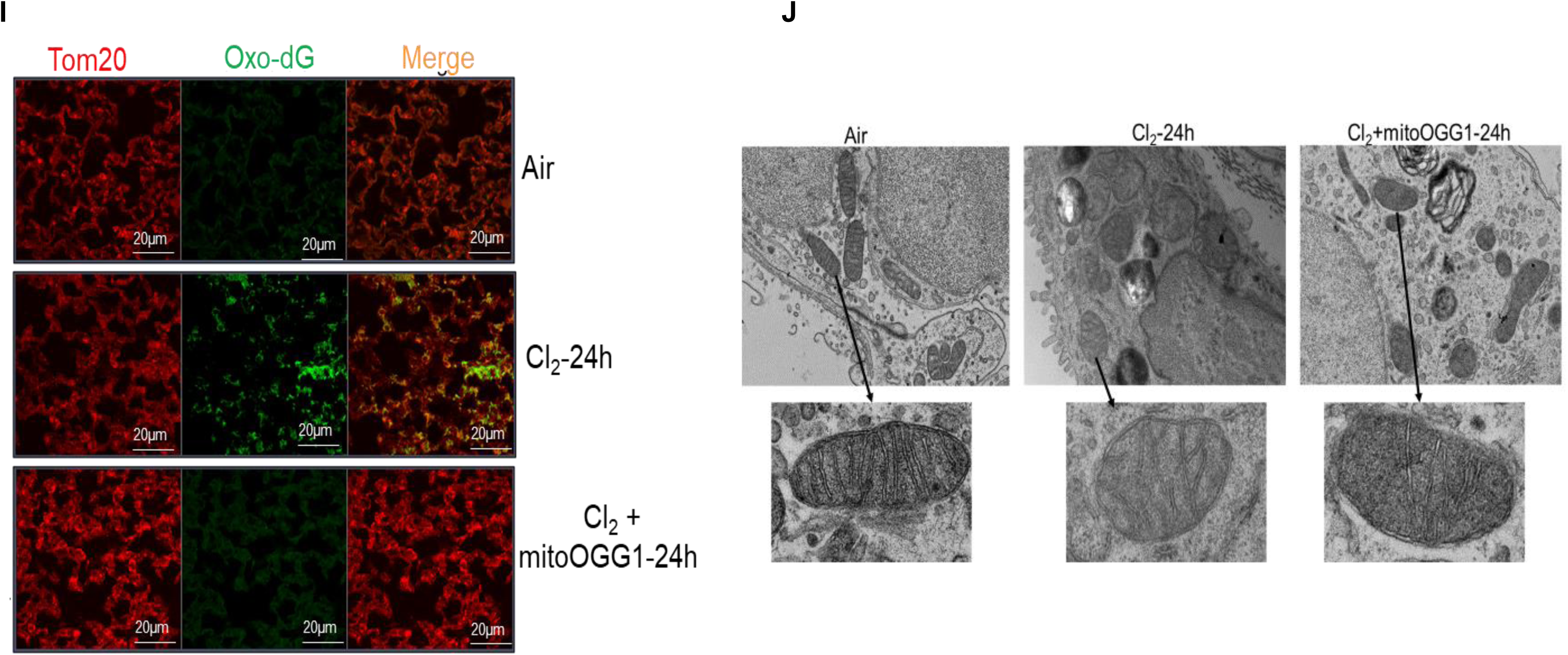
Adult Male and Female (C57BL/6) mice were exposed to Cl_2_ gas (500ppm for 30 min) and returned to room air. One hour later, mitoOGG1 (1 mg/Kg BW in 50 µl buffer) or vehicle were instilled intranasally. Mice were sacrificed and Lung tissues were collected at different time points post Cl_2_ exposure. (**A**) mRNA expression of OGG1 was measured by real-time PCR. Data are shown as fold change after normalization with the housekeeping gene (18s); means±1 SEM, each symbol represents data from a different mouse. (**B**) Immunoblot of lung tissues using antibodies against mouse OGG1 and β-tubulin at the indicated conditions. Each lane represents data from a different mouse. (**C**) Fold changes of lung OGG1 proteins of data shown in **B** at 24 h post exposure. Data are normalized to the corresponding values of β-tubulin. each symbol represents data from a different mouse. (**D**) PCR Amplification of long (∼11kb) and short fragments (∼100bp) of mitochondrial DNA (mtDNA) using genomic DNA isolated from lung tissues from air, post 24hrs of Cl_2_ and following instillation of vehicleor mitoOGG1 (1 mg/kg BW in 50 µl of buffer) at 1 h post exposure. (**E**) Intact mtDNA in lung tissue (expressed as fold change of the air value), assessed by RT-PCRfor the indicated conditions; means±1 SEM, each symbol represents data from a different mouse. (**F)** Intact lung mtDNA injury at 1 h post exposure measured by Southern blots in the mitochondria D-loop in the presence and absence of Fpg; means±1 SEM, each symbol represents data from a different mouse. (**G**) Immunoblot of lung tissues with antibodies against GAPDH (a cytoplasmic marker) and Tom20, an outer mitochondrial membrane protein. Each lane represents data from a different mouse. (**H**) Dot blot showing 8-Oxo-dG DNA lesions in mtDNA, isolated from fractionated mitochondria from lung tissue for the indicated conditions. Each dot represents a single animal. (**I**). Fixed tissues from the right lung, upper lobe of mice at the indicated conditions (Air, post 24hrs of Cl_2_ and Cl_2_ with mitoOGG1),immunostained with antibodies against 8-oxo-dG (Green) and Tom20 (Red). Colocalization of 8-oxo-dG and Tom20 (orange color) indicates the presence of 8-Oxo-dG in the mitochondria at 24 h post Cl_2_. One slide per mouse; n=3 mice for each group. (**J**) Transmission electron microscopy images of mitochondria with abnormal morphology in mice lungs. Damaged mitochondria with matrix swelling and collapsed cristae was observed post Cl_2_ exposure. Scale bar; 1 μm, 600 nm. Data from three slides for one mouse for each group. Data are shown in fold change and SEM; One-way ANOVA followed by the Tukey t-test adjusted for multiple comparisons (PRISM 10).

### Formation of lung mtDNA Lesions and 8-oxoG in mice exposed to Cl_2_ and returned to room air

We then subjected lung tissue to differential centrifugation as described in the methods and isolated the mitochondrial fraction. As shown in **Fig. 1G**, the mitochondrial fraction immunostained with Tom20, a mitochondrial outer membrane marker, while the cytoplasmic fraction immunostained with GAPDH. Dot-blots of the lung mitochondrial fraction with 8-OxoG, showed the presence of 8-OxoG (the most prevalent oxidative modification by reactive oxygen species) in mtDNA (**Fig. 1H**), isolated from the lungs of mice exposed to Cl_2_ and returned to room air at 24 h post exposure. Furthermore, immunofluorescence studies of lung tissues, showed co-localization of 8-OxoG with Tom20, in lung tissues from mice at 24 h post exposure (**Fig. 1I**) but not in those of air controls. We also investigated the changes in subcellular ultrastructure of mice lungs at 24 h post Cl_2_ exposure by using Transmission Electron Microscope (TEM). Mitochondria with swollen matrix and collapsed cristae was accumulated post 24h of Cl_2_ exposure, whereas Intranasal instillation of mitoOGG1 improved the cristae structure (**Fig. 1J**). In contrast, no significant injury to nuclear DNA was detected as shown by unchanged levels of nuclear DNA (detected by real time PCR) and phospho-H2AX, detected by Western blotting at 24 h post exposure (**Supplementary** Fig. 1B, C).

### Intranasal Instillation of mitoOGG1 in mice post Cl_2_ exposure mitigates indices of acute lung injury

We have previously shown that mice exposed to Cl_2_ and returned to room air develop progressive lung injury to both the upper airways and blood gas barrier (2, 5, 8, 33, 37). We thus investigated whether intranasal instillation of mitoOGG1 reverses oxidative injury to lung mtDNA and mitigated the extent of lung injury. We labelled mitoOGG1 with the positron emitter ^89^Zn, as described in the Materials and Methods section. After formulation in 5% ethanol/1X PBS, the 1.85 MBq (50 µCi) of ^89^Zr-mitoOGG1 (**Fig. 2A**) was instilled intranasally in mice at 3 h post exposure to air or Cl_2_ and imaged them at various times post exposure. Large amounts of ^89^Zr-mitoOGG1 were detected in the lungs, heart, liver and kidneys of both air and Cl_2_ exposed mice, shortly after and at 6 h post instillation (**Fig. 2B-C-D**). Small amounts were also present in the lungs 24 h post injection (**Fig. 2D,E**).

**Figure 2.**
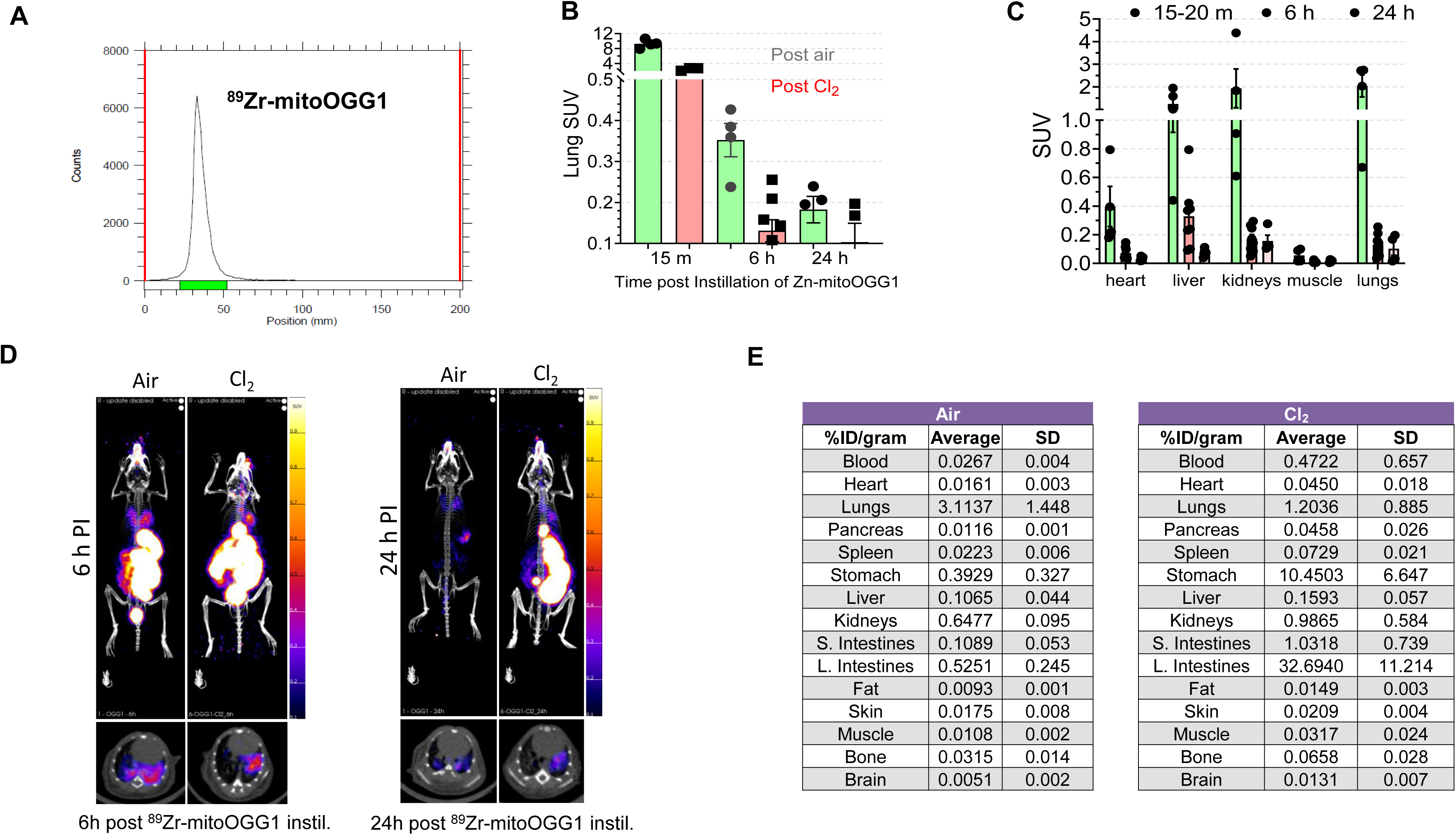
Distribution of intranasally instilled ^89^Zn-mitoOGG1 in air and Cl_2_ exposed mice. Male and female C57BL/6 mice were exposed to air or Cl_2_ (500ppm for 30min) and returned to room air. Three hours later, ^89^Zr-mitoOGG1 (1mg/kg BW in 50 µl of buffer) was instilled intranasally. (**A**) Radio-iTLC of ^89^Zr-OGG1 (R_f_ of ^89^Zr-mitoOGG1 is 0, R_f_ free ^89^Zr ∼ 1). (**B**) Mean Standard Uptake Values (SUV_mean_)±1 SEM (n=4 mice per group) of ^89^Zr-mitoOGG1 levels measured in the lungs of mice in the control (post air) and treated (post Cl_2_ gas exposed) mice over time (**C**) SUV_mean_ comparison of heart, liver, kidney, muscle and lungs at various points post instillation in the Cl_2_ exposed mice. SUV_mean_ values are means ± 1 SEM; each symbol represents an individual mouse. **(D)** Maximum Intensity Projection (MIP) and axial slice PET/CT images of intranasally instilled ^89^Zr-mitoOGG1 at 6 h and 24 h post-instillation in control/air and Cl_2_ exposed mice. **(E)** Biodistribution of ^89^Zr-mitoOGG1 in control/air and Cl_2_ exposed mice at 24 h post instillation. n=4 for each group. Data are expressed at % injected dose/gram (%ID/g), avg = average and SD = standard deviation.

Intratracheal instillation of mitoOGG1 in mice at 1h post Cl_2_ diminished the Cl_2_-induced large increase of airway resistance (**Fig. 3A**) and elastance (**Supplementary** Fig. 2A) following methacholine challenges, the concentration of protein in the BAL (**Fig. 3B and** **Supplementary Fig 2B**), as well as the increase of lung wet/dry weights at 24 h post Cl_2_ exposure (**Fig. 3C**). In addition, H&E sections of lung tissues of mice that were exposed to Cl_2_ and returned to room air at 24 h post exposure showed the characteristic appearance of acute lung injury in contrast to the lung sections of mice that were instilled with mitoOGG1 at 1h post Cl_2_ had almost normal appearance (**Fig. 3D**). At 24 h post exposure, there was a large influx of neutrophils in the BALF and a concomitant small decrease of alveolar macrophages; instillation of mitoOGG1 did not alter cell type composition in the BALF (data not shown).

**Figure 3.**
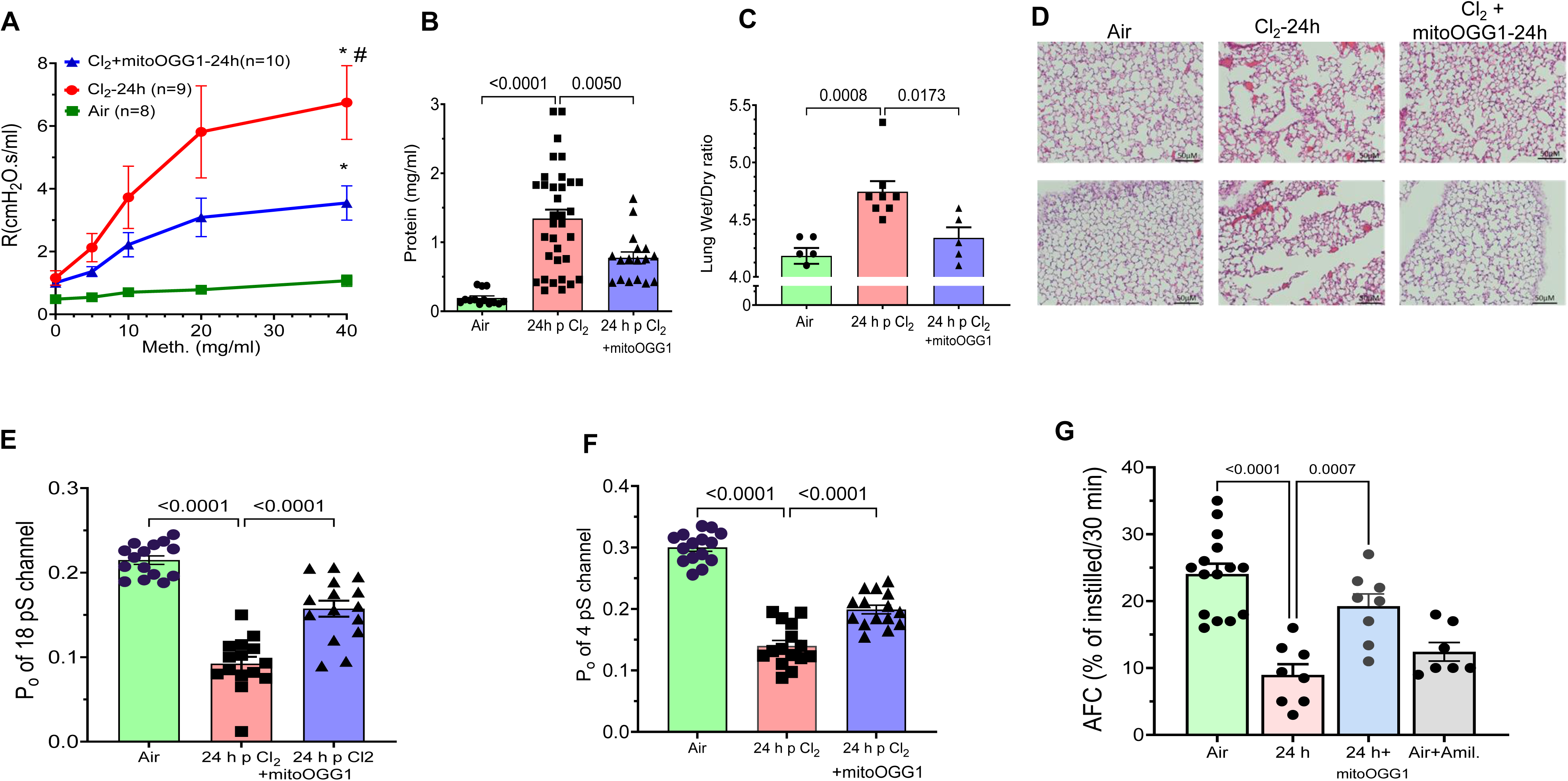
Instillation of mitoOGG1 reduces Cl_2_-induced acute lung. Equal numbers of male and female C57BL/6 mice were exposed to air or Cl_2_ (500 ppm/30 min) and returned to room. They were then instilled with mitoOGG1 (1mg/kg BW in 50 µlin buffer) or Vehicle. Airway Resistance (**A**) was measured by flexiVent as described in the Materials and Methods. Mean values ± 1SEM. (**B**) Concentration of proteins in the broncho-alveolar lavage fluid (BALF) at 24 h in vehicle or mitoOGG1 instilled mice as described above ;means±1 SEM, each symbol represents data from a different mouse. (**C**) Lung W/D weight ratio were elevated at 24 h post Cl_2_ in mice instilled with vehicle but not in mice instilled with mitoOGG1; means±1 SEM, each symbol represents data from a different mouse. (**D**). H&E staining of lung tissue sections shows extensive diffuse injury in airways and distal lung regions that were mitigated by instillation of mitoOGG1. Characteristic appearance in lung sections from right upper lobe, Multiple images were obtained from each mouse (n=3 mice per group) with identical results. (**E, F**). Open probability the amiloride sensitive (4 pS) and cation (18 pS) channels obtained by cell-attached patches of ATII cells in lung slices. Values are mean ± SEM. Two to three patches per mouse; n=3-4 mice per group. Statistical analysis by One-way ANOVA followed by the Tukey t-test adjusted for multiple comparisons (PRISM 10). (**G**). Sodium-dependent alveolar fluid clearance measured in vivo at 24 h post Cl_2_ exposure in mice instilled with saline or mitoOGG1 at 1 h post exposure. Each symbol represents a different mouse. Statistical analysis by one-way ANOVA followed by the Tukey’s posttest.

We then assessed whether instillation of mitoOGG1 mitigated the Cl_2_-induced injury to alveolar type I and type II cells amiloride sensitive sodium (ENaC) and cation channels in slices cut from the lungs of mice at 24 h post exposure. As previously reported(31), exposure of mice to Cl_2_ decreases the open probability of both the 18 pS cation and the 4 pS amiloride-sensitive Na^+^ channels (ENaC) in both ATI and ATII cells patched in the cell attached mode. Instillation of mitoOGG1 at 1h post exposure, increased the open probability of both channels significantly **(Fig. 3 E,F and** **supplementary Fig 2C**). To show the physiological significance of these findings, we measured sodium dependent alveolar fluid clearance (AFC) in vivo, an important property of the alveolar epithelium to decrease alveolar edema when the permeability of the blood gas barrier was increased. As shown in **Fig. 3G**, the alveolar epithelium of mice breathing air clears about 25% of isotonic fluid and this process is significantly decreased in the presence of amiloride, an inhibitor of Na^+^ channels. There was significant decrease of alveolar fluid clearance across the alveolar epithelium of mice post exposure to Cl_2_, the value at 24 h being similar to that following instillation of amiloride in air breathing mice. Instillation of mitoOGG1 at 1h post exposure returned alveolar fluid clearance to almost air control levels (**Fig. 3G**).

### Cl_2_ induced activation of pattern recognition receptors and cytokines is mitigated by mitoOGG1 instillation

Lungs and plasma were harvested from mice 24h post exposure to air or Cl_2_. Total DNA was extracted from equal volume of plasma and BALF and the concentration was measured by nanodrop. There was a significant increase in the amount of DNA in the BAL and plasma from mice injected with vehicle and sacrificed at 24 h post Cl_2_ as compared to air control mice (**Fig. 4A, B**); in contrast, DNA content in the BAL and plasma of mice instilled with mitoOGG1 at 1 h post exposure and sacrificed at 24 h were decreased significantly (**Fig. 4A,B**). RT-PCR analysis of the whole lung lysate demonstrated that at 24 h post Cl_2_, there was a significant increase of lung NLRP3 mRNA (**Fig. 4C**), TLR9 mRNA (**Fig. 4D**) as compared to air controls; in contrast, these variables were at control levels in the lungs of mice instilled with mitoOGG1 (**Fig. 4C, D**). The TLR9 dependent cytokines IL-1β,IL-6 as well as KC/GRO were also higher in the plasma of Cl_2_ exposed animals at 24 h post exposure but instillation of mitoOGG1 returned them to air control values (**Fig. 4E, F, G**). Taken together these results demonstrated that the exposure to Cl_2_ activates mitochondrial DAMP-dependent inflammatory pathways, which may ultimately lead to acute lung injury.

**Figure 4.**
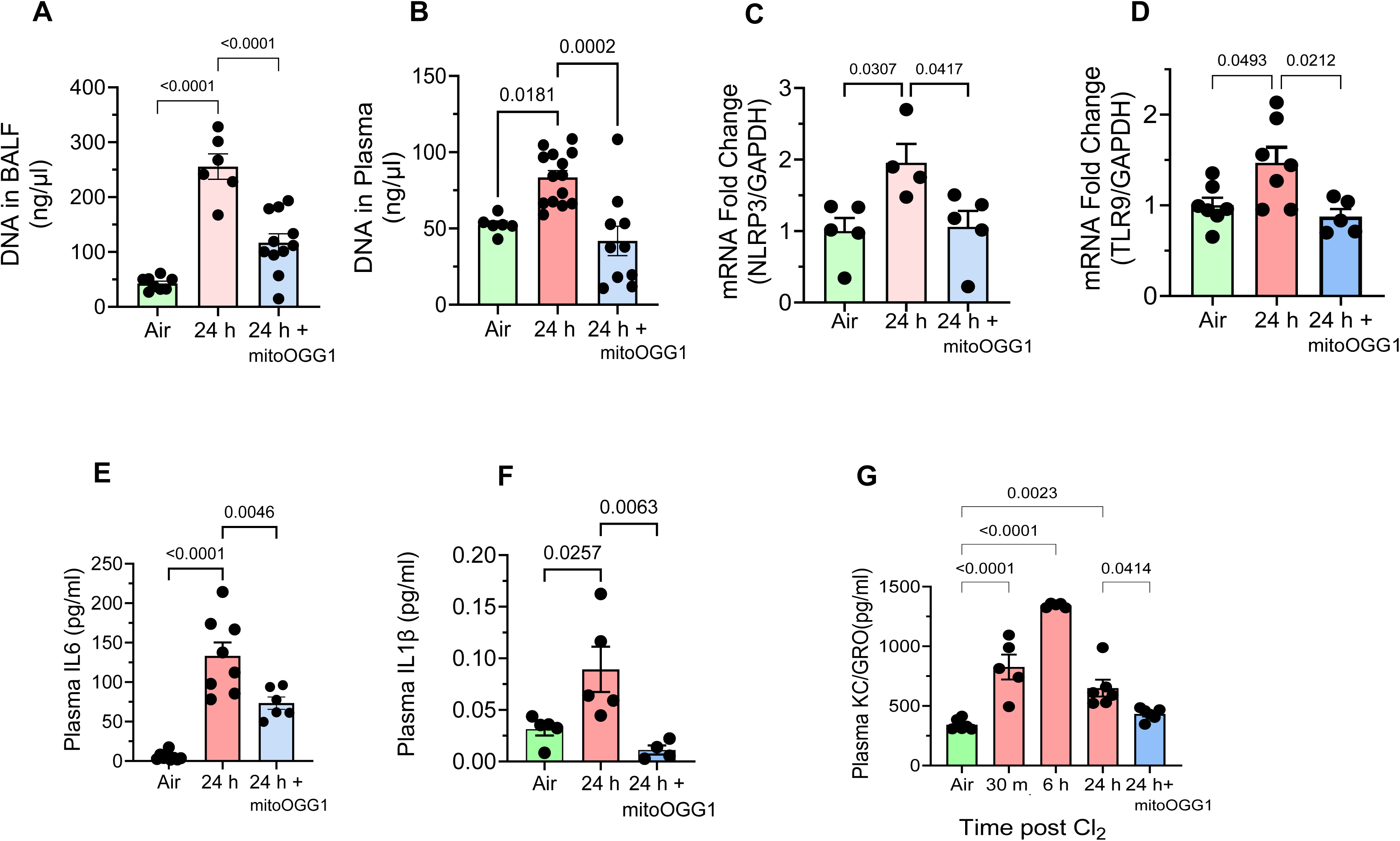
mitoOGG1 fusion protein mitigates Cl_2_-induced lung inflammation. Male and female C57BL/6 mice were exposed to air or Cl_2_ (500ppm, 30min). One hour later, mice were instilled intranasally with mitoOGG1 (1mg/kg BW in 50 µl of sterile saline) or saline, and then lungs were harvested 24hr post Cl_2_ exposure. (**A**) Total DNA in BALF and (**B**) plasma. (**C, D**) mRNA levels of NLRP3,TLR9 in lung tissues. (**E**,**F,G**) Plasma levels of the cytokines (IL6, IL-1β and KC/GRO measured by ELISA immunoassay. Values are mean ± SEM; each symbol represents a different mouse, Statistical analysis by one-way ANOVA followed by the Tukey’s posttest.

### Instillation of mitoOGG1 increases survival and repairs the extent of COPD-type injury

We exposed equal numbers of C57BL/6 male and female mice to 400 ppm Cl_2_ for 30 min and returned them to room air. We chose this regiment to increase number of survivors after one week of exposure. At six h post exposure, mice were instilled intranasally with either mitoOGG1 (0.5 µg/g BW in 50 µl of saline) or saline alone. As shown in **Fig. 5A**, mice receiving mitoOGG1 survived considerably longer that those receiving saline (p=0.02870). At ten days post exposure, approximately 80% of mitoOGG1 instilled mice were alive as compared to about 40% of those receiving saline. Saline instilled mice sacrificed at 14 days post exposure exhibited a leftward shift of the inflation and deflation semi-static pressure volume curves (**Fig. 5B**) and total lung capacity (**Fig. 5C**) which were mitigated by instillation of a single dose of mitoOGG1 at six h post exposure (**Fig. 5B, C**). Transgenic mice with global (mitochondrial and nuclear) OGG1 depletion (OGG1^-/-^) exposed to Cl_2_ exhibited much higher mortality as compared to their control littermates (**Fig. 5D**). In addition, OGG1 ^(-/-)^ mice exposed to 400 ppm for 30 min and returned to room air for 24 h had significantly higher mtDNA injury and protein in the BALF as compared to the corresponding values of their wild-type controls (**Fig. 5 E,F**).

**Figure 5.**
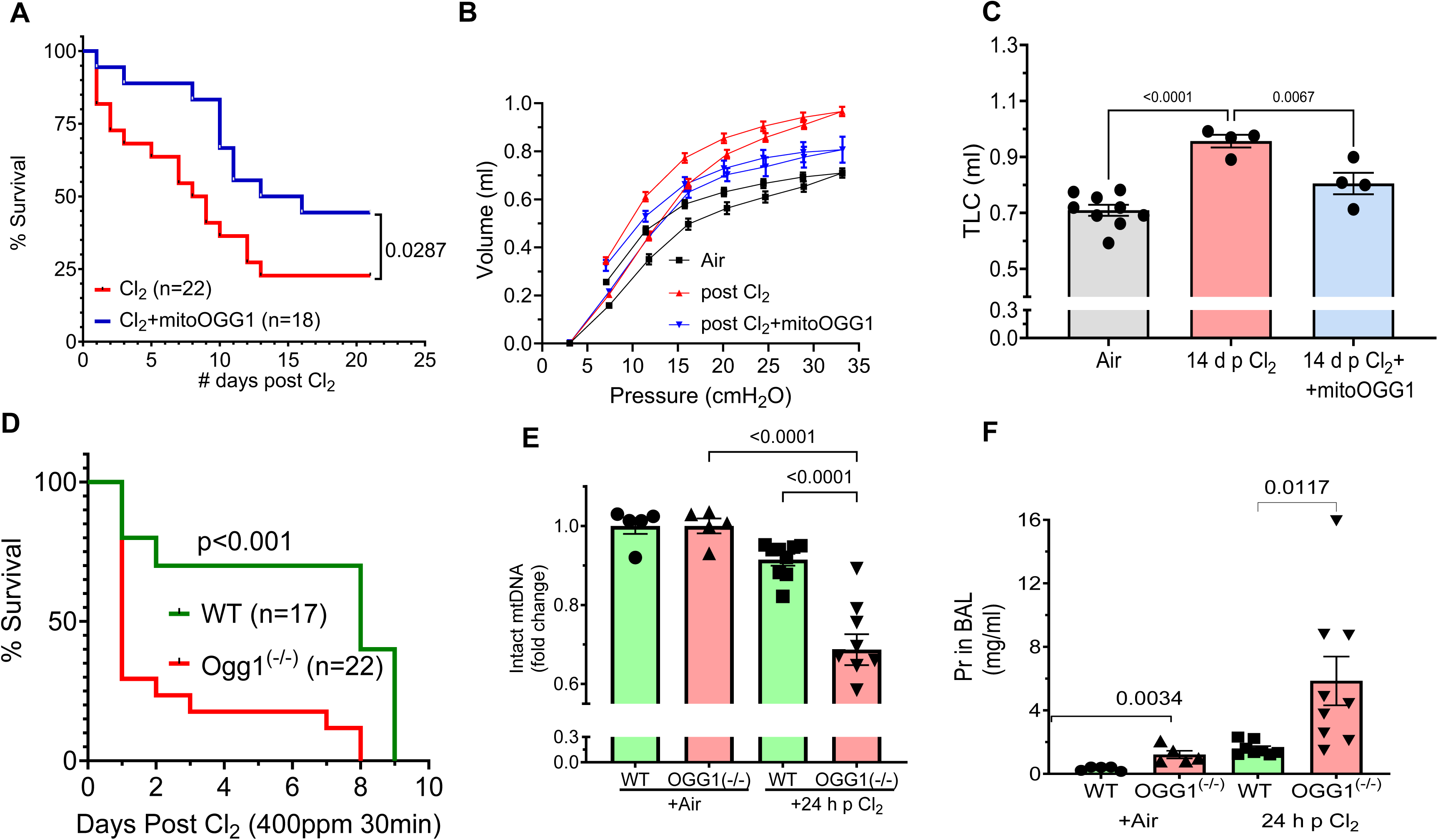
Instillation of mitoOGG1 improved survival and decreased lung injury at 14 d post exposure. Male and Female C57BL/6 mice were exposed to air or Cl_2_ (400 ppm for 30 min) and returned to room air. Post 6hrs of Cl_2_ exposure, mice were intranasally instilled with mitoOGG1 (0.5mg/kg BW in 50 µl of sterile saline) and saline and were kept under observation for 21 days (**A)** Kaplan-Meier curve showing improved survival in intranasally instilled mitoOGG1 mice as compared to the saline group; statistical significance was calculated by using GraphPad PRISM. .(**B**) Quasi-static pressure volume loops measured by flexiVent at 14 days post exposure of male and female C57 mice to air or Cl_2_. Inflation and deflation curves are shown for each group. Lung compliance is measured from the linear portion of the deflation loop. Means ± 1 SEM; n=9 for air; n=5 for Cl_2_ and n=4 for Cl_2_+mitoOGG1 for (**C**). Total lung capacity (TLC) defined as the lung volume at 22 cmH2O pressure for the indicated groups. Values are means ± SEM; n=4-6 for each group. 1-way ANOVA with Tukey post-test. (**D**). Transgenic mice with global (mitochondrial and nuclear) OGG1 depletion (OGG1^-/-^) exposed to Cl_2_ exhibited much higher mortality as compared to their littermate controls in the FBV and C57B/L6 background. Kaplan-Meyer Curves generated by PRISM. Statistical Analysis by the Log-rank (Mantel-Cox) test (PRISM). Numbers indicate the number of mice for Wild-type controls (n=17) and global OGG1(-/-) mice in the same background (mixture of FBV-C57BL/6). (**E-F**). Global OGG1^(-/-)^ and wild-type controls were exposed to 500 ppm for 30 min and returned to room air. At 24 h post exposure, OGG1(^-/-^) mice had lower mtDNA injury (**E**) and BAL protein levels (**F**) as compared to their wild type controls. Values are means ± SEM; Each symbol represents data from a different mouse. 1-way ANOVA with Tukey post-test.

### Cl_2_ induced changes in the lung mitochondrial proteome is mitigated by mitoOGG1

Previously, we reported major proteomics changes in the lungs of mice exposed to 500 ppm Cl_2_ for 30 min and returned to room air for 24 h(37). Herein we opted to study changes in the lung proteome following exposure to a sublethal injury by exposing mice to 500 ppm Cl_2_ for 20 min and compare to the findings with the data obtained at 24 h post 500 ppm Cl_2_/30 min. As shown in **supplementary Fig. 3 A,B** this resulted in significantly less injury to alveolar epithelium, as assessed by the concentration of plasma protein in the BAL, and mtDNA, assessed by RT-PCR as compared to exposures at 500 ppm for 30min. Injury to ATII cell ENaC and cation channels was also significantly less and was repaired completely by instillation of mitoOGG1 (**supplementary Fig. 3C,D**). Thus, these measurements enabled us to assess major lung proteomic changes at 24 h post Cl_2_, and the extent to which they were repaired by mitoOGG1 instillation, prior to the development of mtDNA and the physiological injury to the blood-gas barrier.

Spectral counts for all samples processed for the three different groups: (Air, 24 h post Cl_2_; 24 h post Cl2 instilled with mitoOGG1) for 2287 proteins are shown in **supplementary Table S1.** Non-targeted global proteomics analysis of the lung of mice exposed to 500 ppm Cl_2_ for 20 min and returned to room air for 24 h showed the presence of 1543 proteins for which normalized spectral counts of at least three of the four animals were above zero. (**supplementary Table S2**). From this group, 188 proteins were statistically different from their air control values (86 increased and 102 decreased). Major pathways altered at 24 h post exposure to Cl_2_ (500 ppm/20 min), ordered by -log(p value) and z-score, are shown in **Table 1**. Exposure to Cl_2_ resulted in activation of EIF2 signaling, neutrophil degranulation, acute phase response, inactivation of mitochondrial signaling among others. Instillation of mitoOGG1 at 1 h post Cl_2_ modified 118 lung proteins (48 increased and 70 decreased) at 24 h post exposure as compared to lung of vehicle instilled mice (**supplement Table S3 and Volcano plot Fig. 6A**). The top 39 proteins in this group (i.e., altered by Cl_2_ and reversed by mitoOGG1) were used to construct the heat map shown in **Fig. 6B** and the principal component analysis (**Fig. 6C**). Both the heat map and the PCA analysis show the presence of clear and major differences in the lung proteomes. In addition, the heat map in **Fig. 6D** shows that instillation of mitoOGG1 altered the expression of mitochondrial or mitochondrial associated proteins that were modified at 24 h post Cl_2_.

**Figure 6.**
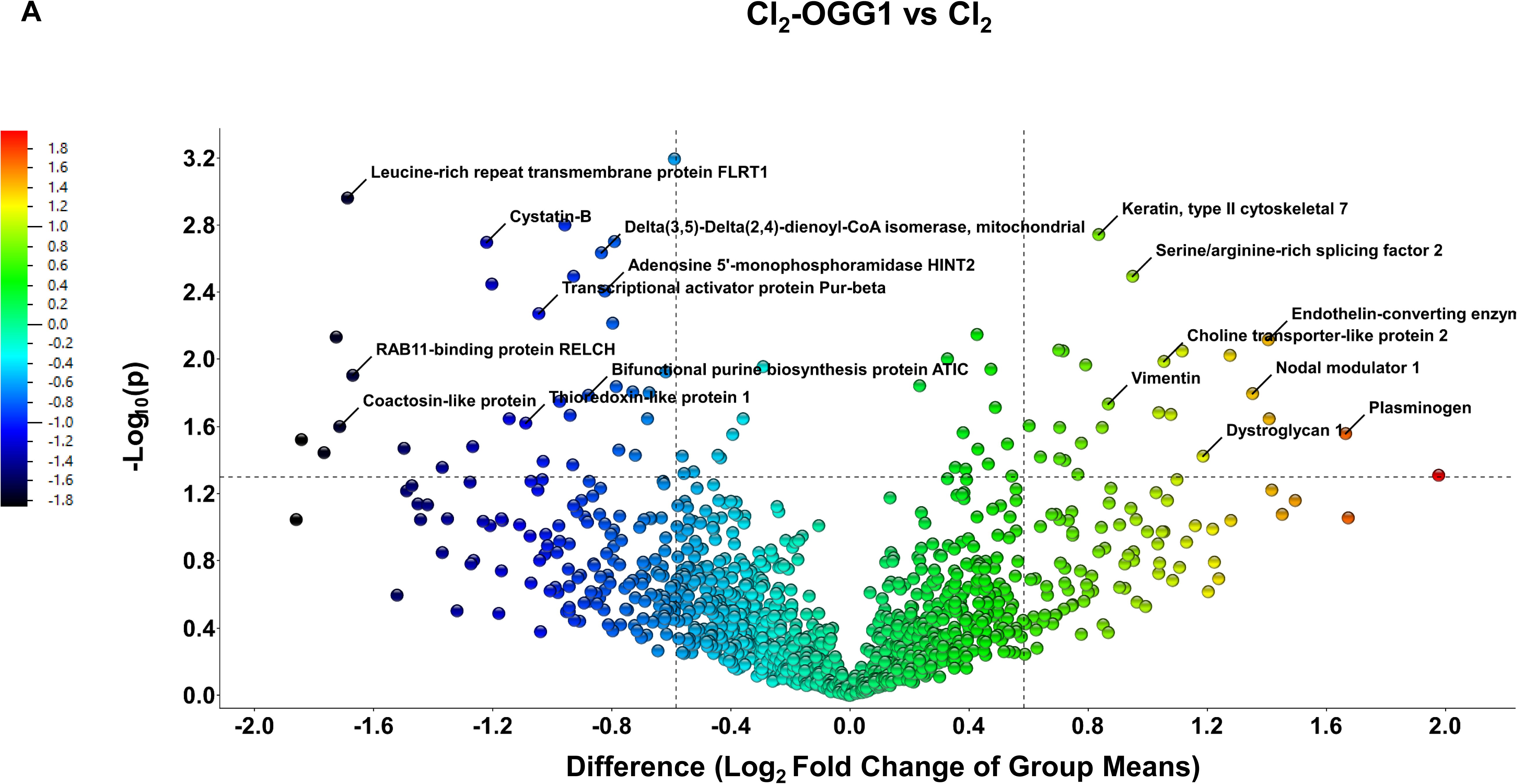
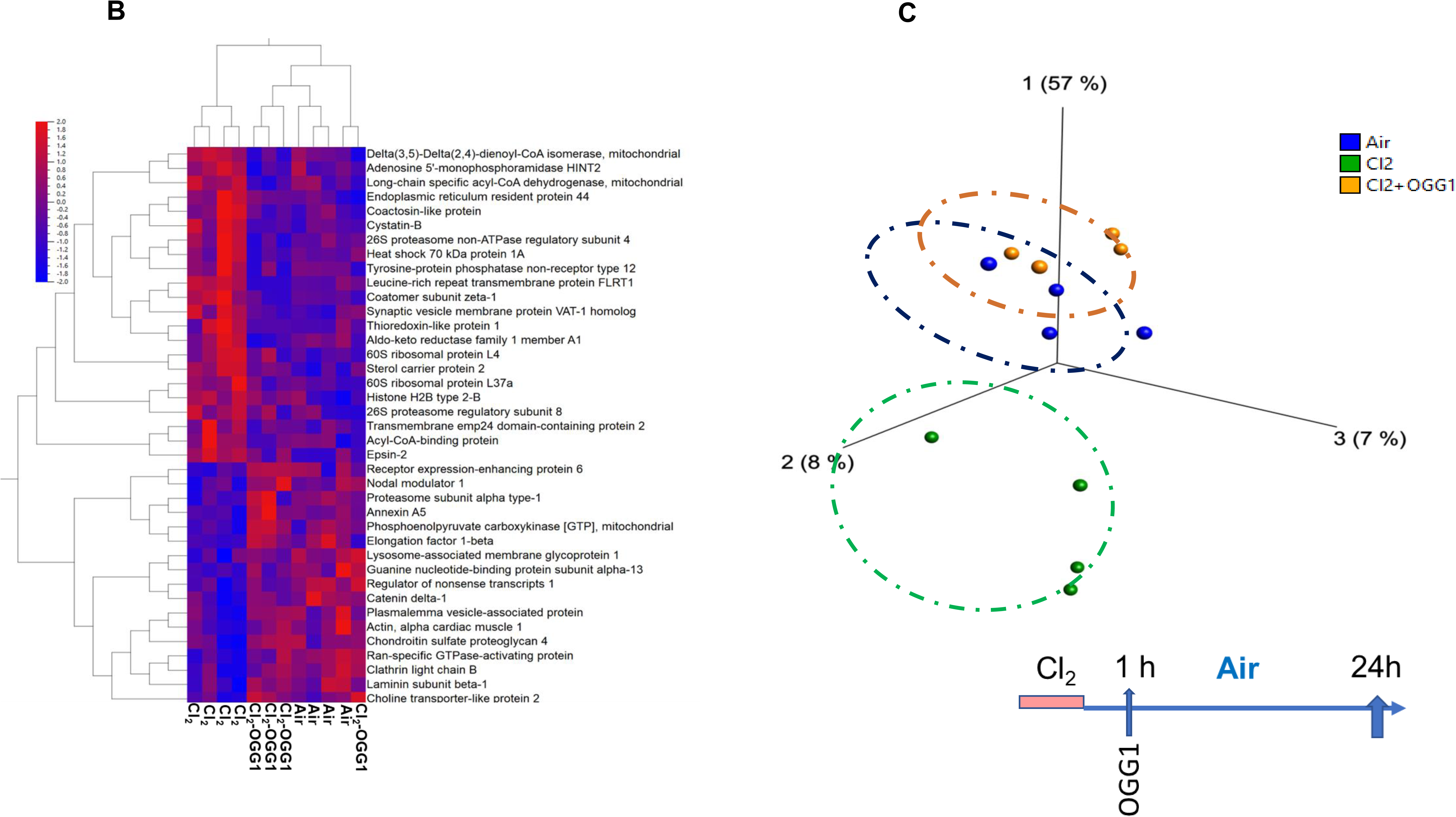
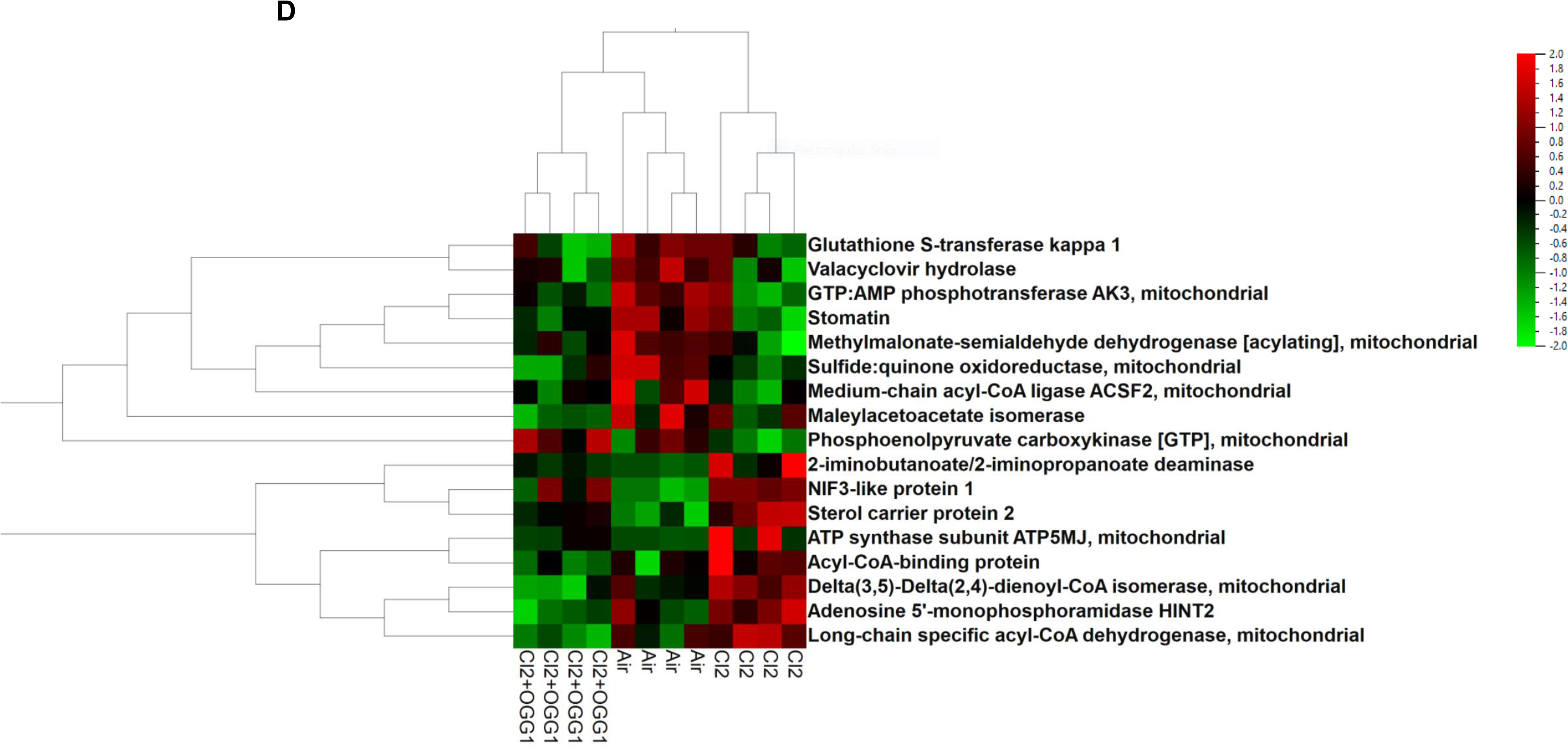
Changes in Lung Proteome post chlorine (Cl_2_) exposure. Male and female C57BL/6 mice were exposed to air or Cl_2_ (500 ppm for 20 min) and returned to room air. Twenty- four h later the lungs were removed and proteins were processed for global proteomics analysis as described in the Materials and Methods. (**A**). Volcano plot of the 181-lung protein changed (increased or decreased) at 24h post Cl_2_ exposure and reversed by mitoOGG1. Statistical analysis by ANOVA in the entire data set. Horizontal dotted line indicates the p=0.05 level of significance. The two vertical dotted lines shows proteins that were increased or decreased by at least 1.5-fold. The horizonal line shows the p=0.05 value (-(log(1.3)). (**B**) Two-dimensional hierarchical analysis heat map of the top 39 proteins. Each column presents data from a mouse for the indicated conditions (**C)** The principal component analysis (PCA) complements the heat map by using a cluster approach that is similar across all animals analyzed. Each dot represents data from a different mouse (**D**) Changes in Lung Mitochondrial proteins for the indicated conditions.

**Table 1.**
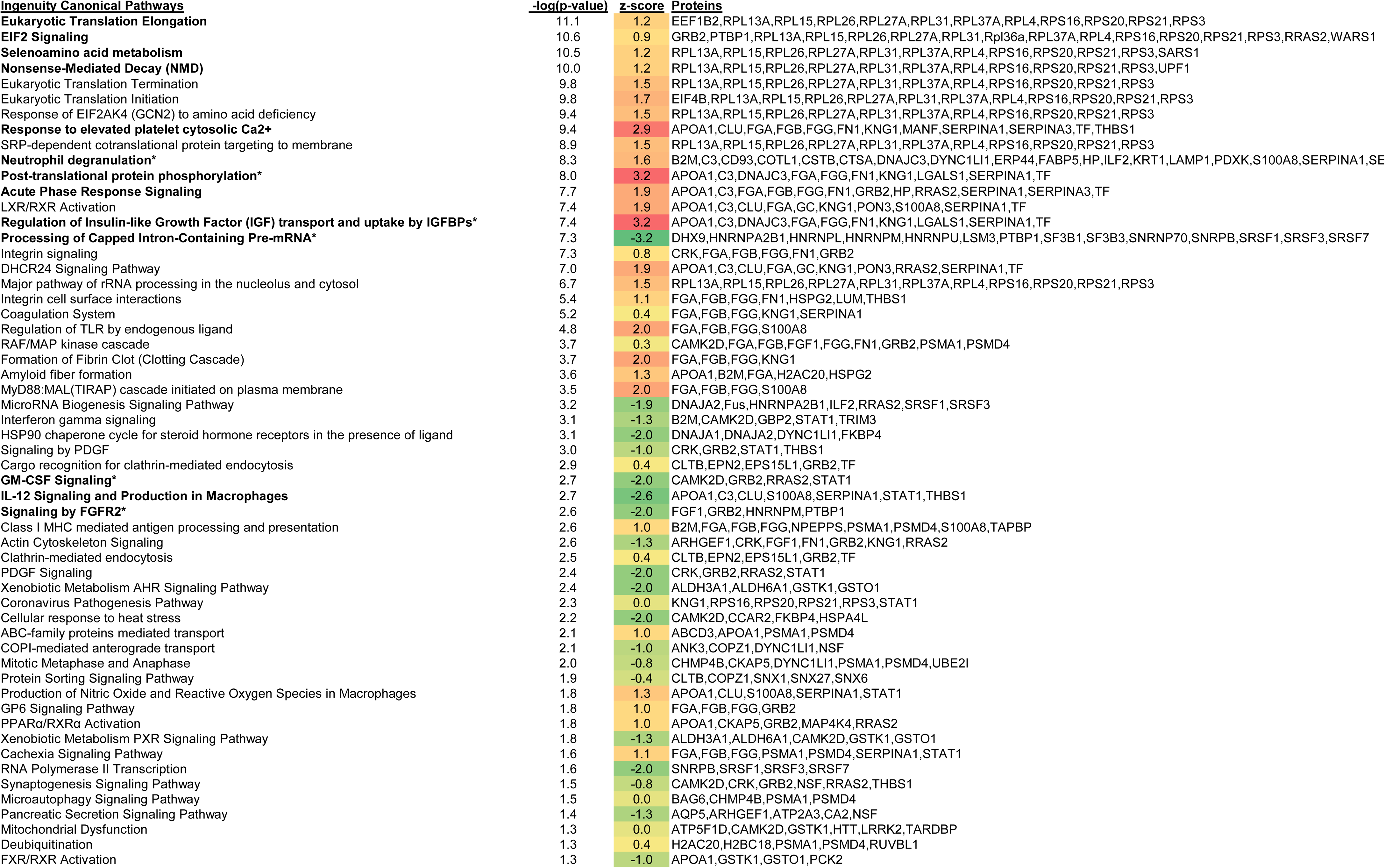
Ingenuity Canonical Pathways modified at 24 h post Cl2 based on non-targeted proteomic analysis of lung proteins shown in Supplemental Table 1. Pathways are ordered by - log(p) and z-scores (predicted pathway activities; orange-red active/green inactive). Proteins associated with each pathway are also shown.

### mitoOGG1 improves the mitochondrial bioenergetics of H441 cells

Human club-like epithelial cells (H441) were grown in air-liquid interface, exposed to Cl_2_ gas at 100 ppm for 10 min and placed in an incubator at 95% air, 5% CO_2_. As shown in the immunoblots of **Fig. 7A and B**, incubation of H441 cells with mitoOGG1 at 1 h post Cl_2_, increased levels of OGG1 mainly in their mitochondria and the levels were decreased significantly post exposure to Cl_2_. Confocal microscopy examination of H441 cells stained with MitoTracker and MitoSox, showed significant colocalization of these two mitochondrial markers at 3 h post Cl_2_ exposure, indicating increased production of mitochondrial reactive species; however, no such colocalization was seen in H441 cells treated with mitoOGG1 (**Fig. 7C**). In the next set of experiments, we added either mitoOGG1 (1 µg per ml) or vehicle at the apical surface of each well at 1 h post exposure to Cl_2_ or air. Oxygen consumption rate was then measured with a SeaHorse following addition of Oligomycin, FCCP and rotenone/ antimycin A. As shown in **Fig. 7 D, E, F, G**, exposure of H441 cells to Cl_2_ decreased basal respiration rate (BRR), maximal respiration rate (MRR) and ATP-dependent respiration; addition of mitoOGG1 at 1 h post exposure returned BRR and MRR and ATP production near to air value.

**Figure 7.**
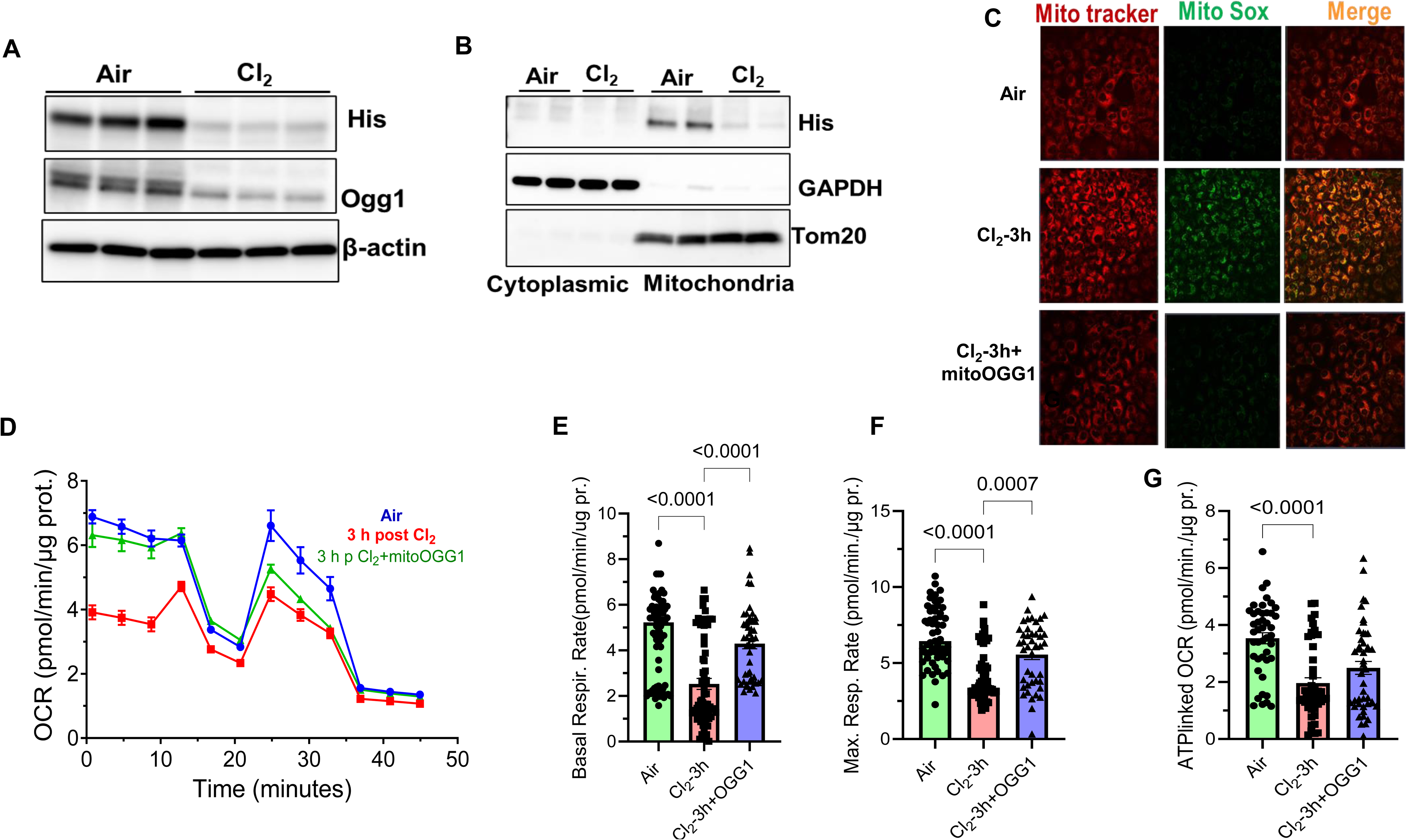
Exposure to Cl_2_ disrupts the mitochondrial function. H441 cells exposed to Cl_2_ (100 ppm for 10 min) and placed in an incubator in 95% air, 5% CO2. Post 1hr of Cl_2_ exposure, the cells were treated with mitoOGG1 (2.5 µg/ml), added in the apical compartment, and measurements of bioenergetics were performed by Sea Horse 3h later. **(A)** Immunoblot showing exogenous and endogenous OGG1 levels detected by anti-His Tag and OGG1 antibodies, respectively. **(B)** Immunoblot showing localization of mitoOgg1 fusion protein in H441 cells, either post Cl2 exposure or in Air, in cytoplasmic and mitochondrial fractions. (**C**) At three hours post exposure, H441 cells were stained with Mito tracker (Red) and Mito SOX (green) and examined by confocal fluorescence microscopy. Notice co-localization of Mito tracker and Mito SOX (yellow) in Cl2 exposed cells indicating the presence of mitochondrial reactive oxygen species. (**D**) Oxygen consumption rates (OCR; pmol/mg/µg protein) of H441 cells exposed to Cl_2,_ returned to 95% air and 5% CO2, transfected with mitoOGG1 or vehicle. Measurements with a SeaHorse at 3 h post exposure. (**E, F, G**) Basal, maximal OCR and ATP linked OCR measurements. Values are mean ± SEM. 1-way ANOVA with Tukey post-test.

## Discussion

The major findings of this study are: (1) Exposure of male and female C57BL/6 mice in concentrations and time intervals likely to be encountered in the vicinity of industrial accidents (27, 51, 52) decreases the concentrations of OGG1 in lung tissues resulting in oxidative injury to lung mitochondria DNA (mtDNA) and acute lung injury reminiscent of ARDS starting tat 6 h post exposure; (2) circulating mtDNA DAMPs activate intracellular cGAS-STING and NLRP3 and the endosomal DAMP receptor, TLR9 resulting in the release of inflammatory cytokines; (3) post exposure instillation of mitoOGG1 in Cl_2_ exposed mice, reaches the airways and distal lung regions; mitoOGG1 instillation initiated mtDNA repair, mitigated the extent of acute ARDS-type injury, seen at 24 h post exposure, the development of COPD-type injury,, seen at 14 days post-exposure and improved post Cl_2_ survival; (4) exposure of C57BL/6 mice to Cl_2_ results at 24 h post exposure in extensive modification of the lung mitochondrial proteome which is repaired to a large extent by mitoOGG1 instillation; (5) transgenic mice with global (mitochondrial and nuclear) OGG1 depletion (OGG1^-/-^) experience much higher mortality post exposure to Cl_2_ as compared to their wild type controls and (6) exposure of human airway Club-like cells to Cl_2_ results in compromised mitochondrial bioenergetics which is repaired by post exposure administration of mitoOGG1. Taken as whole these in vivo and in vitro studies provide convincing evidence that damage to lung mtDNA is a key contributor to the propagation of acute and chronic injury post-exposure to Cl_2_.

Previously, we have shown the presence of increased levels of heme, a major prooxidant, in the plasma of both mice and humans post exposure to Cl_2_ (5, 37) and that removing free heme by injections of hemopexin prevents injury to the mitochondria. OGG1 is a key enzyme that plays an important role in repairing oxidative damage to mtDNA. We found that the expression of OGG1 in lung tissue was significantly downregulated post Cl_2_, which led to increased mtDNA damage. Previous studies have shown that depletion of endothelial cell OGG1 potentiated ROS-induced mtDNA damage and increased apoptosis (17). Furthermore, HeLa cells defective in mitochondrial OGG1 activity were prone to oxidative mtDNA damage, which was reversed by overexpression of OGG1(44). We and others have shown that administration of mitoOGG1 suppress and reverse oxidative mtDNA damage and physiologic dysfunction in multiple rodent models of inflammatory injury characterized by ROS stress and mtDNA damage(14, 24, 25, 29, 35, 48). In agreement with previous studies, we found that mtDNA is more prone to damage than nuclear DNA due to its lack of protective proteins and proximity to mitochondrial ROS sites. (23,9). Similarly, we found that Cl_2_ exposure of C57BL/6 mice and H441 cells leads to increased mitochondrial ROS production resulting in mitochondrial DNA damage and dysfunction whereas Intranasal delivery of mitoOGG1 in mice reduced mtDNA damagen and 8-oxoG lesions and helped retain the lung mitochondrial cristae structure,

To confirm the intranasal delivery of mitoOGG1 to the lungs of air and Cl_2_-exposed mice, we used PET/CT imaging by radiolabeling recombinant the mitoOGG1 protein with ^89^Zn. The majority of instilled mitoOGG1 was localized in the lung and some traces were also found in systemic organs. Notably, previous studies have also reported that traces of instilled recombinant proteins were identified in other organs were due to their mucociliary clearance in the trachea. These particles mixed with salivary secretions and are swallowed, entering the gastrointestinal tract for excretion through the bowel (47).

Repairing mitochondrial oxidative injury by a single intranasal instillation of mitoOGG1 partially repaired Cl_2_-induced injury to the upper airways (as shown by decreased airway reactivity in response to methacholine) and damage to the blood gas barrier, as shown by decreased levels of plasma proteins in the BAL and improved alveolar structure in H&E staining. In addition, we reported for the first time that mitoOGG1 reversed injury to lung epithelial sodium channels, a key pathway, in limiting the accumulation of fluid in the alveolar epithelium when the integrity of the blood-gas barrier was compromised. We thus, explored the molecular mechanisms and assessed how intranasal instillation of mitoOGG1 mitigates lung injury and improves its function.

While it is well known that mtDNA damage is directly cytotoxic, accumulating evidence shows that mtDNA damage-dependent DAMP formation also plays a critical pathogenic role by activating downstream inflammatory cascades that could propagate injury to remote sites, even if they are unaffected by the initial insult (45, 49, 57). This may be critical for the delayed effects of Cl_2_ gas where the inflammatory pathways activated by mtDNA-DAMPs are engaged in a feed-forward inflammatory pathways that could propagate injury despite control of the initiating stimuli (29, 36, 38, 41, 42). Observational studies in human patients show that plasma mtDNA- DAMPs are biomarkers of clinical outcomes in disparate disorders, including cardiometabolic diseases, sepsis, ischemia-reperfusion injury, trauma, and others(23, 30, 40, 46, 48). Experiments in laboratory animals or cell culture models show that exogenous mtDNA DAMPs recapitulate many aspects of disease by activating pro-inflammatory responses(12, 29, 56, 57). Exogenous mtDNA DAMPs also cause oxidative damage to the endogenous mitochondrial genome, thus activating a feed-forward cycle of mtDNA damage and mtDNA DAMP formation(29). Pro-inflammatory actions of mtDNA DAMPs are linked to stimulation of nucleic acid receptors at the heart of the innate immune response: the intracellular cGAS-STING and NLRP3 and the endosomal DAMP receptor, TLR9(6, 10, 45, 55). In our study, we found that the inflammatory pathways were activated upon exposure to Cl_2_ gas and mito-OGG1 reversed these processes. These findings suggest that the instillation of mitoOGG1 reduced the activation of NLRP3 and TLR9, resulting in reduced inflammation and improved lung injury.

The chronic effects of chlorine inhalation and treatment by mitoOGG1 are important to understand as our research ensures that the restorative effects we have observed translate to measurable improvements in mortality and morbidity. Mice exposed to chlorine saw a significant increase in mortality with ∼20% of mice dying 1-day post-exposure and 50% dying 10 days post-exposure. However, when chlorine-exposed mice were given mitoOGG1 6h post-exposure, we saw a decrease in mortality at all time points, with only 5% mortality at day 1 and 35% mortality at day 10. Additionally, 80% of mice receiving no treatment died by the end of the study, while our experimental mitoOGG1 group saw only 55% mortality at day 21. We tested our hypothesis in a global deletion of OGG1 (^-/-^) knockout mice and found much higher mortality than their wild-type controls post-Cl_2_ exposure. These data highlight the crucial role of OGG1 in repairing and mitigating lung injury and decreasing mortality post-exposure to Cl_2_.

Previously, our unsupervised global proteomics and systems biology studies, coupled with Western blotting studies, showed an increase of phosphorylated E2Fa, in the lungs of mice at 24 h post exposure to 500 ppm Cl_2_ for 30 min. (37). The fact that hemopexin, the major scavenger of free heme, a major oxidant released in the circulation following fracture of red blood cells, indicates that that activation of E2Fa was the result of increased oxidant stress, as we demonstrated herein. Activation of E2Fa has been linked to mitochondrial dysfunction and the onset of apoptosis(9). In this study we show that instillation of mitoOGG1 reversed the Cl_2_ induced modification of a large number of lung tissue as well as mitochondrial-associated and mitochondrial proteins, such as Annexin A5, Choline transporter-like protein 2, Ran-specific GTPase activating protein, Long-Chain specific acetyl-CoA dehydrogenase, ATP synthase subunit ATP-5MJ, NUIF3-like protein 1 and others (see Figure 6) which regulates the diverse biological functions and pathways in the mitochondria or nucleus including ATP synthesis,mitochondrial function, cell proliferation, transcription, replication, inflammation and DNA repair (11, 18).

Mitochondria are responsible for generating ATP which is the fuel for all metabolic functions. Herein, we found that treatment of H441 cells with mitoOGG1 improved mitochondrial function compared to the vehicle post exposure to Cl_2_. Moreover, Cl_2_ leads to increased ROS production and downregulation of OGG1 expression, but the mitoOGG1 mitigates the damage and restores the bioenergetics and mitochondrial function.

Based on our findings, we propose that Cl_2_ exposure leads to ROS production in lung tissue, which results in mtDNA damage due to a lack of protective proteins. Downregulation of OGG1 leads to increased inflammation and mtDNA lesions. As a result, reduced mitochondrial function and acute/chronic lung injury result in death. Intranasal Instillation of mitoOGG1 protein reversed the phenomenon by reducing the Cl_2_-induced lung injury with improved survival.

## Author contributions

S.M., J.M. and S.D. conceived and designed the research. S.D., Z.Y., E.M.S., A.L. I.A., S.A., V.M.P., M.I.H,J.L.B, S.R.F. performed the experiments. S.M., S.D., Z.Y., T.J., S.R.F., and J.A.M. helped with analyzing data. S.M.,S.D., A.L., M.G., S.E.L, S.A., T.J. J.A.M., M.G. interpreted results. S.D. drafted the manuscript; S.M. edited the manuscript. J.M performed the proteomics measurements and systems biology analysis. All authors discussed at each stage of the manuscript and have approved the final version of the manuscript.

## Materials & Correspondence

Correspondence and material requests can be addressed to Dr. Sadis Matalon.

## Materials and Methods

### Animals

Wildtype (C57BL/6) 8-12 weeks old male and female mice (∼20-25 g, body weight) were purchased from Charles River Laboratories (Wilmington, MA). After arrival, the mice were kept in mice facility for at least 72h of acclimatization before using them for experiments. Breeding pairs of global OGG1(-/-) mice in the FBV-C57BL/6 background and wild type controls were provided by Dr. Gillespie at the University of South Alabama, All experimental protocols on mice were approved by Institutional Animal Care and Use Committee of University of Alabama at Birmingham (IACUC-UAB) under APN 22366, 22606 and 22835.

### Exposure to Cl_2_

Before Cl_2_ exposure, intraperitoneal long-acting buprenorphine (1 mg/Kg, to minimize anxiety and discomfort) was injected into all unanesthetized mice according to UAB IACUC guidelines. Mice were then placed inside cylindrical chambers located in an environmental hood inside a negative pressure room and exposed to either Cl_2_ gas (500 or 400 ppm for 20-30min) as described previously(5, 8, 33) (34). At the end of the exposure, the mice were returned to their cages and observed during the day at least every 2 h and once during the night. They were allowed access to food and water ad libitum. For the survival studies, mice were monitored for 2 wk. to determine the level of mortality between each group. Mice were sacrificed if their body weight dropped below 30% of their initial weight following IACUC approved protocol.

### Generation and administration of recombinant mitoOGG1

The generation of constructs was done as described previously. Briefly, Codon optimized constructs were placed in plasmids for expression in E. coli of fusion proteins containing EndoIII coupled to a TAT sequence to facilitate cellular uptake, the MTS from MnSOD, a hemaglutin (HA) tag for immunological localization and a histidine tail as previously described (24). All reagents for fusion protein production were obtained from Sigma-Aldrich (St. Louis, MO, USA) unless otherwise indicated.

MitoOGG1 (1mg/Kg or 0.5 mg/Kg in 50 µl of sterile saline) was administered in the external nares of conscious mice at 1 or 6 h post exposure to Cl_2._

### Measurements of plasma proteins in bronchoalveolar lavage fluid (BALF)

Mice were anesthetized with an I.P. injection of ketamine and xylazine (100 and 10 mg/kg body weight, respectively) and their lungs were lavaged with 1 ml of PBS; ∼0.7–0.9 ml was recovered in all groups (7, 37). Recovered aliquots of lavage fluid were kept on ice and centrifuged immediately at 3000 xg for 10 min to pellet the cells. Supernatants were removed and stored on ice and the protein concentrations were measured with the BCA Protein Assay Kit (Cat. No 23225; Thermo Fisher Scientific, Rockford, IL).

### Measurements of Respiratory Mechanics by flexivent

**C57**B/l6 mice were exposed to air or Cl_2_ (500ppm or 400ppm for 20- 30min) and 1hour later, mitoOGG1 was instilled intranasally (1mg/Kg of BW in 50 µl saline). The respiratory mechanics was measured by using the forced oscillation technique (flexiVent, SCIREQ) as described previously(32, 50). Briefly, at 24 h post exposure, the mice were anesthetized with pentobarbital sodium (50 mg/kg ip.; Vortech Pharmaceuticals, Dearborn, MI) and paralyzed with pancuronium (4 mg/kg ip; Gensia Sicor Pharmaceuticals, Irvine, CA), intubated, then connected, and mechanically ventilated by a flexiVent FX system (SCIREQ, Montreal, QC, Canada) as described previously (2). Lungs were challenged by increased concentrations of aerosolized methacholine (0 to 40 mg/mL) applied within 10 s of the start of each measurement. Airway responses were recorded every 15 s for a consecutive 3 min following the aerosolization of methacholine and airway resistance and elastance were calculated according to manufacturer’s instructions.

### Lung wet-to-dry weight ratio measurements

C57B/l6 mice were exposed to air or Cl_2_ (500 or 400ppm - for 20- 30min) and 1hour later, mitoOGG1 was instilled intranasally (1mg/Kg of BW in 50 µl saline in each nostril). At 24 h post exposure, the mice were deep anesthetized with an intraperitoneal injection of ketamine and xylazine . The chest was cut opened, and the lungs were removed. Lungs were pre-weighed and placed in an oven for drying at 55°C for 7 days. Post 7 days the lungs were re-weighed and the wet-to-dry weight ratio was calculated.

### Alveolar Fluid Clearance (AFC) measurements

C57B/l6 mice were exposed to air or Cl_2_ (500- for 20- 30min) and 1hour later, mitoOGG1 was instilled intranasally (1mg/Kg of BW in 50 µl saline in each nostril). AFC was measured at 24 h post exposure as described previously(5, 32). **Briefly,** Mice were anesthetized with an intraperitoneal injection of Ketamine (80- 100mg/kg) and xylazine (5-10mg/kg). The trachea was exposed and cannulated with an 18- gauge intravenous catheter trimmed to 0.5 inch long . Mice were positioned in the left decubitus position, and 0.5ml of 5% BSA was instilled over 30sec and infused with 0.1ml air in the catheter to clear the dead space and position the fluid in the alveolar spaces. Mice were ventilated with a flexiVent as described previously(5, 32) At the end, the remaining alveolar fluid was collected using a 1ml syringe and protein concentrations were measured with the BCA method as described above. AFC expressed as a percentage of the instilled volume, was calculated using the following formula: AFC = (1- Ci/C30)/0.95, where Ci and C30 are the protein concentrations at times zero and 30min, respectively.

### Patch clamp in ATII cells in lung slices

Alveolar type I and type II cells in lung slices were patched in the cell attached mode as described previously (5, 32, 34). Briefly, C57B/l6 mice were exposed to air or Cl_2_ (500 or 400ppm for 20- 30min) and 1hour later, mitoOGG1 was instilled intranasally (1 µg/g of BW in 50 µl saline in each nostril). Mice were euthanized using a lethal dose of ketamine and xylazine, then their chests were cut open, the lungs were perfused with Krebs-Ringer solution containing (in mM): 140 NaCl, 3 KCl, 2.5 CaCl_2_, 1 MgCl, 10 glucose, and 10 HEPES (pH ≈7.35–7.4), and subsequently filled with low temperature melting agar through an incision in the trachea to stiffen the tissue for slicing with a microtome (Precision Instruments. Greenville, NC). The lower lobe of the right lung was dissected and mounted on the microtome and sliced into 250μm thick slices. The slices were incubated in DMEM culture media at 37°C in humidified atmosphere of 5% CO_2_ for three hours at which point a slice was transferred to the recording chamber filled with a solution of the following composition (in mM): 130 K-gluconate, 2 MgCl_2_, 10 KCl, 10 glucose, and 10 HEPES (pH 7.4, KOH) and held in place with an anchor (Warner Instruments). The recording pipette resistance ranged from 5-6MΩ when filled with a solution of the following ionic composition (in mM): 140 NaCl, 3 KCl, 2.5 CaCl_2_, 1 MgCl, 5 glucose, and 10 HEPES (pH = 7.4). The recording chamber was mounted onto the stage of an upright Olympus microscope EX51WI (Olympus, Center Valley, PA). ATII cells within the slices were visualized by the accumulation of LysoTracker green (1 μM, ThermoFisher) in the lamellar bodies of ATII cells. Data were recorded and stored onto the hard drive of a computer equipped with pClamp software (Molecular Devices, San Jose, CA) and interfaced to an Axon amplifier Axopatch 200B (molecular devices) with Digitata 1440A (Molecular devices). Data were analyzed using Clampfit (molecular devices) and the open probabilities of sodium channels were calculated as described previously(31).

### MitoOGG1 conjugation and radiolabeling

mitoOGG1 was conjugated to DFO-Bn-NCS as described previously(19). Briefly, to a microcentrifuge tube containing 100 µg (approx. 2.4 nmol) of mitoOGG1, 100 µL of 1M HEPES buffer pH 7.1 and 10 µL of 1 M Sodium carbonate buffer was added DFO-Bn-NCS as a 4-fold molar excess (7.2 µg or 9.6 nmol) dissolved into DMSO. The vial was briefly vortexed then incubated at 37°C for 1 h. After incubation, the conjugate was purified via a Zeba spin column (7 kDa MWCO) pre-equilibrated in 1 M HEPES buffer at pH 7.1. The protein concentration of the solution was determined via a BCA assay using manufacturer’s instructions and was calculated to be 215 µg/mL.

^89^Zr-oxalate (13 MBq or 350 µCi) was produced as previously described(43) neutralized via an equal volume addition of 1 M HEPES, pH 7.1 and a 1/10 addition of 5 M NaOH. A 5 µg aliquot of DFO-Bn-OGG1 was added to the ^89^Zr and the pH was retested and readjusted using 2 M HCl to a pH between 7.0 – 7.4. The reaction was incubated at 37°oC for 45 min and progress assessed checked via iTLC-SG using 50 mM DTPA as running buffer. The final product was adjusted to a concentration of 1 µCi/µL with 5% ethanol/1X PBS buffer.

### PET imaging and post-PET biodistribution

Mice were anesthetized with 2% isoflurane in oxygen and given ^89^Zr-DFO-mitoOGG1 (1 mg/kg of body weight, a total of 50 µL in 5% ethanol/1X PBS buffer) via intranasal instillation (in both the nostrils equally). Mice were imaged at 0 h (20 min dynamic PET, 3 min CT), 6 h (20 min static PET, 3 min CT), and 24 h (25 min static PET, 3 min CT) post instillation on a Sofie GNEXT PET/CT scanner (Sofie Biosciences, Culver City, CA). Immediately after the 24 h image acquisition, mice were euthanized and organs of interest were collected. The radioactivity and weight of tissues were measured on a HIDEX AMG automated gamma counter (Hidex, Turku, Finland). Uptake of radioactivity was calculated as percent injected dose per gram of tissue (%ID/g).

PET images were reconstructed using a 3D-OSEM (Ordered Subset Expectation Maximization) algorithm (24 subsets and 3 iterations), with random, attenuation, and decay correction, and CT images were reconstructed using a Modified Feldkamp Algorithm. Reconstructed PET and CT images were imported into VivoQuant software (Invicro, Needham, MA) and standard uptake values (SUVs) were determined for select tissues (heart, liver, kidneys, muscle, lungs) by hand-drawing regions of interest (ROIs) using CT anatomical guidelines.

### Southern Blot

We used a PCR-based assay to detect oxidized guanine in short sequences of mtDNA D-loop region(1), reported previously to be sensitive to oxidative base modification(2) and to be the source of proinflammatory mtDNA DAMPs activating NLRP3(3). Total DNA was isolated immediately after treatment using a DNeasy Blood and Tissue Kit (Qiagen GmbH, Hamburg, Germany). All samples were eluted with 100 ul of TE buffer. Samples so isolated were treated with formamidopyrminine DNA glycosylase (Fpg) or its vehicle. For Fpg treatment, 250 ng of DNA with 8 units of Fpg in 1× NEBuffer 1 (10 mM Bis-Tris propane-HCl, 10 mM MgCl_2_, 1 mM DTT, pH 7.0) and 100 μg/ml BSA in a volume of 50 μl at 37°C for 1 h. Fpg was then inactivated by heating at 60°C for 5 min. An aliquot containing 10 ng DNA was then used for the PCR assay to detect Fpg-sensitive cleavage sites. PCR primers were designed with Beacon Designer 8.2 software (PREMIER Biosoft International, Palo Alto, CA) and were as follows: D-loop, forward 5’- TCTACCATCCTCCGTGAA -3’, reverse 5’- GCCCTGAAGTAAGAACCA -3’. A standard curve was created using a PCR product from the whole D-loop region. Data are presented as the fraction of intact mtDNA, calculated as the quotient of signal intensities in Fpg- treated and untreated DNA

### Lung Mitochondrial DNA concentrations and DNA lesion Assay

The total DNA was extracted using 25 mg of lung tissue from the mix of left and right lobe, by using QIAamp DNA mini kit, (cat# 51306, Qiagen, Germany). DNA concentration was measured using nanodrop and equal amount of DNA (∼2-5ng) used to perform qPCR as described previously (46). Briefly,to measure the mitochondrial DNA concentration and DNA lesions, we designed two sets of mitochondrial specific primer targeting 11kb of mitochondrial DNA. In the first step, DNA was amplified using the 11kb mitochondrial DNA specific primers, that were designed in-house: Mito-Large (F): CTT TAC GAG CCG TAG CCC AA; -Mito-Large (R): AGC GAA TCG GGT CAA GG. In the second step, Real-time was performed by taking long PCR product as a template, to quantify the intact Mitochondrial DNA concentration with a short mitochondrial DNA specific PCR primer. Mito-Short-(F): GGC CCA TTA AAC TTG GGG GT; Mito-Short-(R): TTA AGG GGA ACG TAT GGG CG. A different set of experiments were also performed to measure the mitochondrial DNA lesions by using Mouse Real-Time PCR Mitochondrial DNA Damage Analysis Kit from Detroit R&D (Cat. No. DD2M; Detroit, MI) following manufacturer’s instructions.

### Cell culture and treatment

Human NCI-H441 cells (Human Lung epithelial cells, Catt No. HTB-174™) were purchased from ATCC (American Type Culture Collection, USA) and revived in FBS supplemented Dulbecco’s Modified Eagle’s Medium (DMEM, Life Technologies, Grand Island, NY) along with antibiotic Penicillin-Streptomycin (Life Technologies, USA) in a humidified incubator at 37°C with 5% CO_2_. Cells were further used by plating them onto 6-well, 12-well, or 10-cm culture dishes and exposed to cl_2_ and air for different time points.

### Subcellular fractionation

H441 were exposed to Cl_2_ (100ppm/10min.) and 24 hours later, cytoplasmic and mitochondrial proteins were fractionated using Mitochondria Isolation kit (Cat No 89874, ThermoFisher Scientific) following the manufacturer’s instructions. Protease and phosphatase inhibitor was added to the extraction reagents before use. Subcellular Fractions were resolved on denaturing SDS-PAGE gels, and fractions purity was confirmed by immunoblotting against GAPDH (cytoplasmic marker, Cat No. 60004-1-Ig, Proteintech)) and Tom20 (Mitochondrial marker Cat No. 11802-1-AP, Proteintech).

### Dot blot analysis

Total DNA was isolated from lungs of mice post exposure Cl_2_ 24h and air using QIAamp DNA mini kit, (cat# 51306, Qiagen, Germany)according to the manufacturers’ instructions. DNA concentration was quantified using Nanodrop and equal amounts of DNA was crosslinked onto positive charge Hybond-N discs (Amersham Hybond-N membranes, cat# RPN82N; Cytiva). To remove unbound DNA, the membrane was quickly washed and blocked using 5% non-fat dry milk. Primary antibody against 8OxoG (1:500, cat# ab48508, Abcam, USA) were diluted in 2% BSA and membrane were incubated for overnight at 4°C followed by HRP-conjugated secondary anti-rabbit antibody (1:5000, cat# 1706515, Biorad, USA) incubation for 1hr at room temperature. After washing the membrane the images were acquired by using ECL (Cat No. 34076, Supersignal West Dura Extended duration substrate, Thermo scientific) in Genesyn imager (Syngene, USA) . Further, Methylene blue solution (Cat No. # 03978, Sigma Aldrich) used to stain membrane to use as a loading control and signal intensity of each dots was analyzed, which has been shown in fold change.

### Protein extraction and Western blot analysis

Total protein was extracted from 25mg of lung tissue by using 300µl of Tissue Protein Extraction Reagent (T-PER; PIERCE cat #78510, Rockford, IL) with a protease and phosphatase inhibitor cocktail (Thermo Scientific, cat # 78442, Rockford, IL), and centrifuged at 12,000xg for 5 minutes to pellet cells and tissue debris. The supernatant was collected and the protein concentration was measured with the BCA protein Assay (PIERCE; cat # 23228, Rockford, IL). Subsequently, 50 µg of protein sample was heated to 95° C for 6 min, cooled on ice and equal volume was loaded on 4-12% Midi-PROTEAN TGX stain-free protein gels (cat# 4568093, Bio-Rad Laboratories); the gel was then transferred to PVDF membranes using the Trans-Blot Turbo Transfer System(BIO-RAD; Hercules, CA), and blocked with 5% fat free milk in 1xTBST(TBST; J77500.K2; ThermoFisher) with tween at room temperature for 1h. After membrane wash, the blots were incubated with OGG1 (Cat No. 15125-1-AP, Proteintech), GAPDH (Cat No. 60004-1-Ig, Proteintech), and β-Tubulin (Cat No. 10094-1-AP, Proteintech), NDUFS1 (Cat No. 60153, CST), TRXR1 (Cat No. 15140S, CST), β-Actin (Cat No. 66009-1-Ig, Proteintech), 6*His-Tag (Cat No. 66005-1-Ig, Proteintech), OGG1 (NB100-106, Novus) in 5% BSA, 1xTBS, 0.1% Tween 20 at 4°C with gentle shaking, for overnight at 4°C. After washing, the membrane was incubated with HRP-conjugated secondary antibodies against mouse (Cat No. 1721011, Bio-Rad Hercules, CA) and rabbit (Cat No. 1706515, Bio-Rad, Hercules, CA) at room temperature for 1 hour. Then, membranes were incubated with Supersignal West Dura Extended duration substrate (Cat No. 34076, Thermo scientific) and placed in a Syngene Imager for visualization of the protein bands. Protein bands were quantified with GENE TOOL Software.

### RNA isolation and quantitative qRT-PCR

Total RNA was isolated from cells and lung tissue using a Qiagen RNA extraction kit (cat# 74106, Qiagen) and the RNA concentration was quantified using nanodrop. 1µg of RNA was reverse transcribed using Revert Aid First Strand cDNA Synthesis Kit (cat# K1691, ThermoFisher Scientific). Real-time PCR (qPCR) was performed to quantify mRNA expression of different gene using gene-specific primers (in a Quant Studio 3 system (Applied Biosystems, Thermo Fisher Scientific) using SYBR green master mix (PowerUp™ cat# A25778, Thermo Fisher Scientific) as per manufacturer’s instructions. Expression of the target genes was normalized with respective Ct value of housekeeping gene (GAPDH or 18S rRNA). All the primers used in this study are listed in **Table S4; supplement**. Gene expression was represented as fold change with respective to the control group.

### Measurements of Reactive Oxygen Species

H441 cells were seeded and were for 24h. Cells were exposed to Cl_2_ (100ppm/10min) and returned to incubator at 37°C in 5% CO2 and post 1h they were treated with mitoOGG1 (1µg/ml) or vehicle (Buffer used to dilute mitoOGG1) for 2hrs. Afterwards, cells were treated with MitoSOX (Cat No. 36009, Thermo Fisher Scientific) and Mito tracker (Cat No. M22425, Thermo fisher Scientific) for 30 minutes at 37°C, after incubation they were rinsed three times with PBS before fluorescent measurements. Cellular ROS was imaged by confocal microscopy (Zeiss TE200-C1).

### Seahorse Assay

Mitochondrial respiration assay was performed using Extracellular Flux Analyzer (Agilent Technologies, Santa Clara, California). To measure Basal and Maximal OCR, H441 cells were seeded and were incubated in incubator at 37°C with 5% CO_2_ for 24h. Cells were exposed to Cl_2_ (100ppm/10min) and returned to incubator at 37°C, 5% CO_2_ for 1 h, post exposure 1h, few wells were treated with recombinant mitoOGG1 (2.5 ug/ml) and Vehicle (Buffer used to dilute mitoOGG1) for 2hrs . Post 3h of Cl_2_ exposure Oxygen Consumption Rates (OCR) were measured, followed by the injection of a series of mitochondrial inhibitors and reagents: [10 μM DCCD (N,N0-dicyclohexylcarbodimide, an ATP synthase inhibitor (Sigma-Aldrich, St. Louis, MI, USA), 10 μM CCCP (carbonyl cyanide 3-chlorophenol hydrazone, a protonophore (Sigma-Aldrich, St. Louis, MI, USA)], 20 μM rotenone [Complex I inhibitor (Sigma-Aldrich, St. Louis, MI, USA)] and either 10 μM antimycin A [Complex III inhibitor (Sigma-Aldrich, St. Louis, MI, USA)] or 1.5 mM BHAM [benzohydroxamic acid, alternative oxidase (AOX) inhibitor (Sigma-Aldrich, St. Louis, MI, USA)]. Measurement cycles consisting of 3 min of mixing, 2 min wait, and 3 min measurement time were completed before and after each sequentially added drug, at least two cycles per condition. The measurements were then used to calculate Basal and maximal OCR.

### Immunofluorescent cell Assay (IFC)

Immunofluorescence was performed by first deparaffinizing on the paraffin-embedded slides of mouse lungs by heating slides at 90°C for 15min. Next, slides were placed for 5min each in a series of dilutions of Xylene and ethanol in separate Coplin jars as follows: 100% xylene, 100% xylene, 50% xylene 50% ethanol. Rehydration of slides was performed using the following dilutions for 5min each: 100% ethanol, 100% ethanol, 95% ethanol, 75% ethanol, 50% ethanol, de-ionized (DI) water. Antigen retrieval was performed by placing slides in an unmasking solution of Tris EDTA 9.0pH and heating for 20min in a sealed rice cooker. The slides were allowed to cool for 10min in DI water before a hydrophobic barrier was drawn around the tissues and Vector Laboratories Bloxall (SP-6000- 100) was added for 15min at room temperature in a wet chamber. The slides were then washed in TBS+0.025% Triton for 5min. A blocking solution created using 2.5% goat normal serum in 1% BSA made in 0.025% TBST was added to the slides for 2hr at room temperature in a wet chamber. After the blocking step, slides were washed in 0.025% TBST three times for 2min each before adding Abcam Goat F(ab) Anti-Mouse IgG H&L (ab6668) 1:1000 for 1hr in wet chamber at room temperature. Slides were quickly rinsed in 0.025% TBST. Primary antibody incubation was performed using 8-hydroxy-2’-deoxyguanosine (Cat No. ab208670, Abcam) at 1:500 and Tom20 (Cat No.11802-1-AP, Proteintech) at 1:200 made in blocking solution and allowed to incubate overnight at 4°C in a wet chamber. Slides were washed two times for 5min in 0.0025% TBST before Alexa Fluor 488 (mouse) and 594 (rabbit) secondary antibodies were added at a 1:500 dilution to incubate at room temperature for 1hr The slides were shielded from light to complete the next steps. Samples were washed three times for 5min in 0.025% TBST and one time for 3min in TBS. Slides were allowed to dry before mounting with Invitrogen ProLong Glass Antifade Mountant with NucBlue (P36985). Slides were stored at 4°C until use.

### Transmission Electron Microscopy

All measurements were conducted in the UAB Electron Microscopy facility. Mouse lungs were intratracheally inflated with 1% osmium tetroxide in 0.1M cacodylate buffer (Caco buffer) and left in 1% osmium overnight. Lungs were cut into 1 mm pieces and rinsed thrice with Caco buffer (each wash 10 min). Afterwards, the cut tissues were added to 1% low molecular weight tannic acid, sonicated and filtered in 0.1M caco buffer for 20min at room temperature and washed thrice with Caco buffer (each wash 10 min), following 2 times wash with deionized water. Tissues were then sequentially dehydrated thrice in Graded Ethanol (50%, 70%, 95%, 100%) and then washed with propylene oxide twice for 10 minutes. Infiltration was done in 1:1 propylene oxide and embedded in 812 resins for overnight. 100% fresh resin was used for 2-3hrs and finally embedded in Embed12 at 60°C in oven for 48hrs.Ultrathin sections were cut using a Leica UC7 ultramicrotome. Thin sections were post stained with Reynold’s lead citrate and ethanolic uranyl acetate. All samples were imaged on a TEM (JEOL 1400 HC Flash electron microscope with an AMT digital camera 100-CX, Jeol). The Electron Microscopy was performed at our core : UAB High Resolution Imaging facility.

### Global proteomics and systems biology analysis

Preparation of samples and proteomics analysis was carried out as described previously by our group (1, 37, 39). After sacrificing the mice, the lungs were perfused and lavaged with saline (0.9% Saline solution) with a catheter inserted in the right ventricle till they were free of blood. Proteins were reduced with Dichlorodiphenyltrichloroethane (DTT), denatured at 70°C for 10 min prior to loading onto Novex NuPAGE 10% Bis-Tris Protein gels (Invitrogen, Cat # NP0315BOX) and separated ∼half way (15min at 200V constant). Peptide digests (8µL each) were injected onto a 1260 Infinity nHPLC stack (Agilent Technologies, Santa Clara, CA). This system runs in-line with a Thermo Orbitrap Velos Pro hybrid mass spectrometer equipped with a nano-electrospray source (Thermo Fisher Scientific, Watham, MA) and all data sets were collected in Collision-Induced Dissociation (CID) mode. The list of peptide identifications generated based on SEQUEST (Thermo Fisher Scientific) search were filtered using Scaffold (Protein Sciences, Portland Oregon) using filters to retain only high confidence IDs while also generating normalized spectral counts (NSC’s) across all samples for the purpose of relative quantification. Relative quantification across experiments were then performed via spectral counting, and when relevant spectral count abundances were then normalized between samples.

**Statistical and Systems Biology Analysis** was performed as described previously (39). For the proteomic data generated, two separate non-parametric statistical analyses were performed between each pair-wise comparison. These non-parametric analyses include 1) the calculation of weight values by significance analysis of microarray (SAM; cut off >|0.6|combined with 2) t-test (single tail, unequal variance, cut off of p < 0.05), which then were sorted according to the highest statistical relevance in each comparison. Gene ontology assignments and pathway analysis were carried out using MetaCore (GeneGO Inc., St. Joseph, MI). Interactions identified within MetaCore are manually correlated using full text articles.

## Statistical Analysis

All statistical analysis were performed using GraphPad (Prism version 8, GraphPad Software, San Diego, CA). The meanL±LSEM was calculated in all experiments, and statistical significance was determined by either the one-way or the two-way ANOVA. For one-way ANOVA analyses, Tukey’s multiple comparisons test was employed, while for two-way ANOVA analyses, Bonferroni posttests were used. Overall survival rate was analyzed using the Kaplan–Meier method. Differences in survival were tested for statistical significance by the log-rank test. A value of pL<L0.05 was considered significant.

---

**Supplementary Figure 1:**
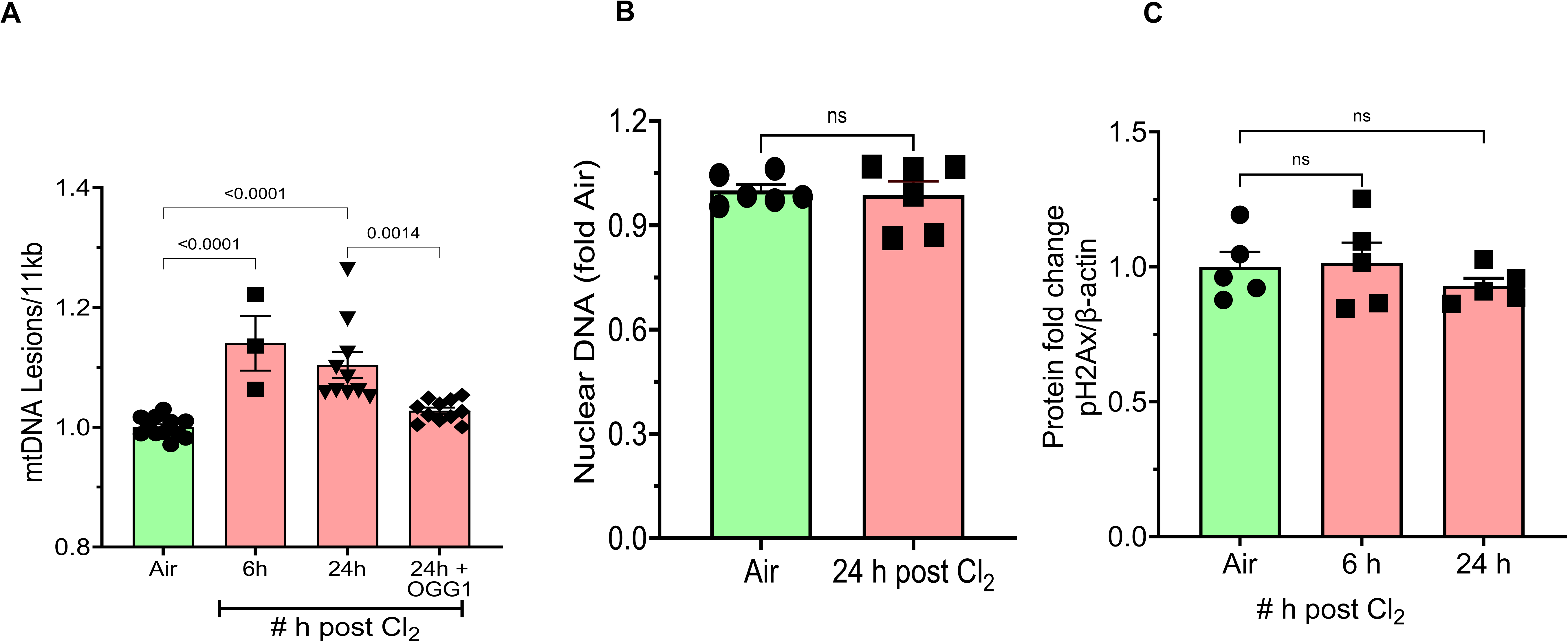
Differences in mtDNA and nuclear DNA lesions post Cl_2_. Adult Male and Female (C57BL/6) mice were exposed to Cl_2_ gas (500ppm for 30 min) and returned to room air. One hour later, mitoOGG1 or vehicle were instilled intranasally (1 mg/Kg BW in 50 µl buffer) or Vehicle. Mice were sacrificed and Lung tissues were collected at different time points post Cl_2_ exposure. (**A**) mtDNA lesions were calculated using the Poisson’s distribution as described in the Methods. Statistical significance was determined by 1-way ANNOVA (PL>L0.05) followed by the Tukey post hoc-test. **(B**) Nuclear DNA injury was assessed by RT-PCR using Nuclear specific set of primers designed in-house. Lung tissues from male and female C57BL/6 mice at the indicated conditions. Means ± 1 SEM; each symbol represents data from a different mouse. **(C)** Densitometric analysis of Immunoblot after normalization with Housekeeping gene. Protein fold change of pH2AX was analyzed by using β-Actin levels. Values are means ± 1 SEM; each symbol represents data from a different mouse. Statistical significance was determined by 1-way ANOVA (PL<L0.05) followed by the Tukey post hoc-test

**Supplementary Figure 2:**
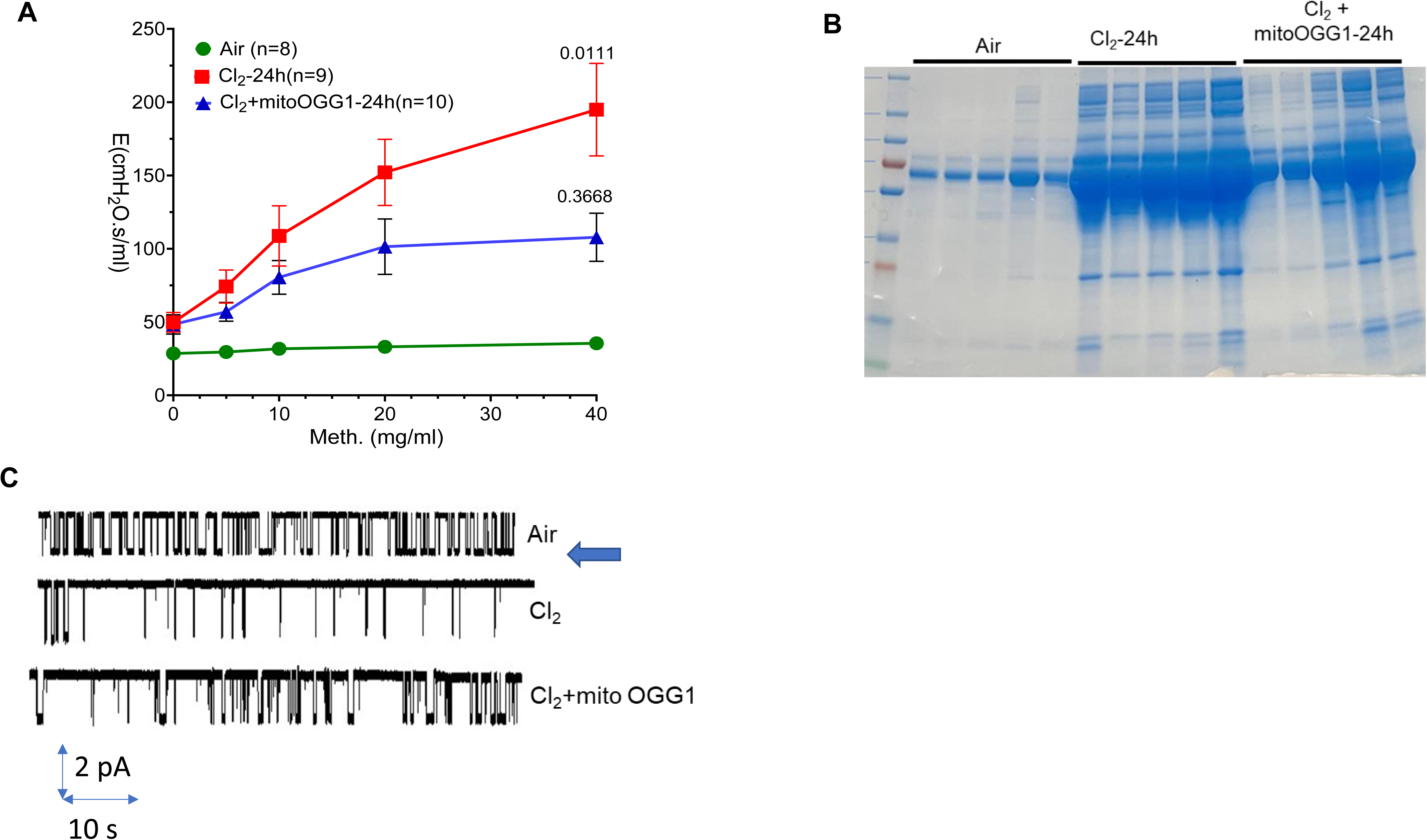
Equal numbers of male and female C57Bl/6 mice were exposed to air or Cl_2_ (500 ppm/30 min) and returned to room. They were then instilled with mitoOGG1 (1mg/kg BW in 50 µl of sterile saline) or Vehicle and Airway Elastance (**A**) by flexiVent as described in the Materials and Methods at given conditions. Values are means ± 1 SEM. (**B**) SDS-PAGE of proteins in the broncho-alveolar lavage fluid (BALF) at the indicated conditions. **(D).** In situ recordings from inside-out patches of the amiloride sensitive Na^+^ channel (ENaC) activity obtained from ATII cells in a lung slice for the indicated conditions. The membrane potential across the patch was −100 mV. The arrow indicates open condition of the channel.

**Supplementary Figure 3.**
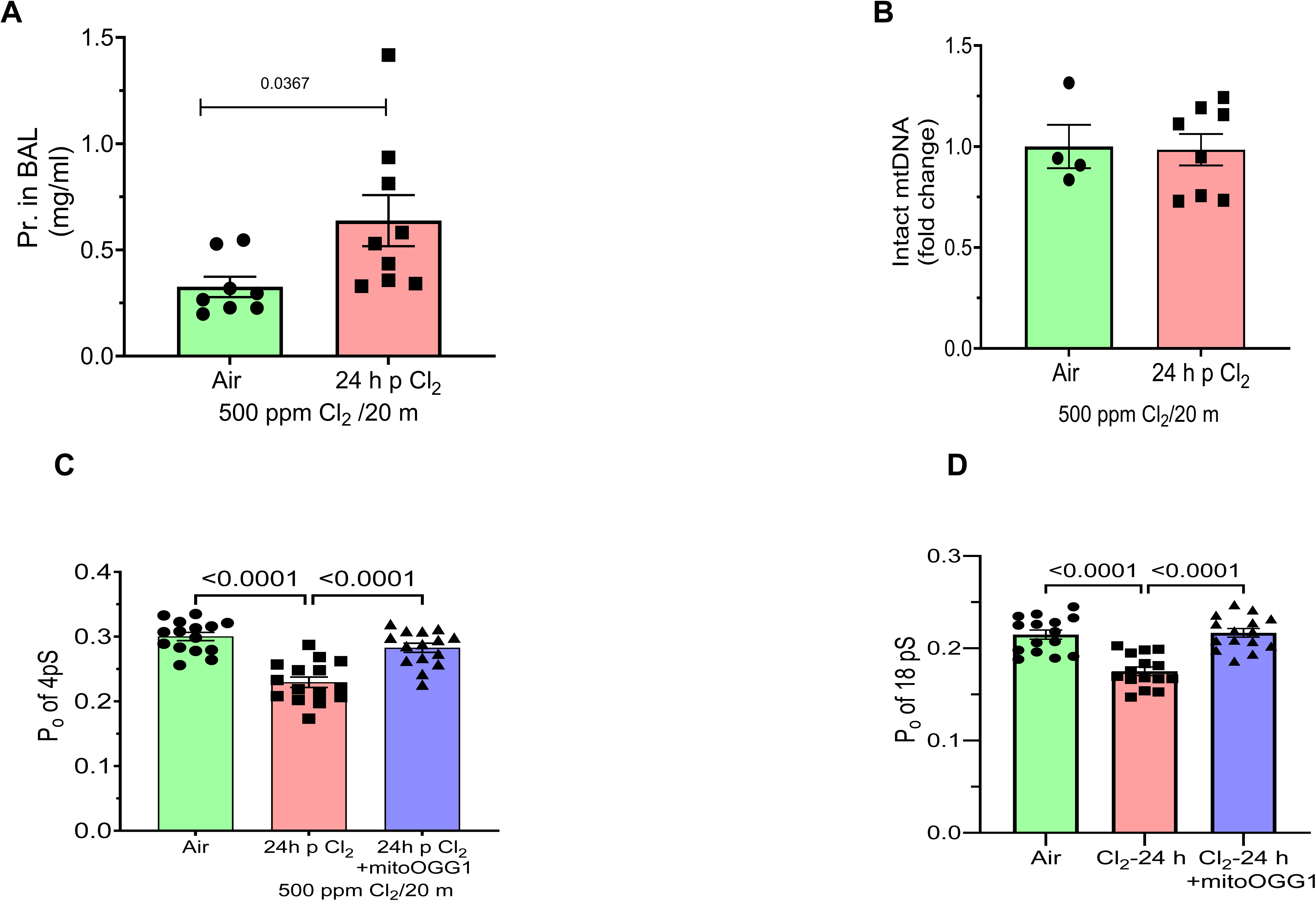
Equal numbers of male and female C57Bl/6 mice were exposed to air or Cl_2_ (500 ppm/20 min) and returned to room. They were sacrificed twenty four h later and the concentrations of proteins in the BAL (**A**) and intact mtDNA in lung tissue (expressed as fold change of the air value), assessed by RT-PCR) (**B**) for the indicated conditions were measured as described in the methods. (**C-D**). Equal numbers of male and female C57Bl/6 mice were exposed to air or Cl_2_ (500 ppm/20 min) and returned to room. At one h post exposure they were instilled with mitoOGG1 (1 mg/kg BW) or saline and were sacrificed 24 h later. ATII cells in lung slices were patched in the cell attached mode and the open probabilities of the 4 pS (ENaC) and the 18 ps cation channels were measured as described in the methods. Values are means ± 1 SEM; values from 2-3 patches from3-4 mice in each group. Statistical significance was determined by 1-way ANOVA (PL<L0.05) followed by the Tukey post hoc-test .

**Supplemental Table 1** Total Spectral Counts (TSCs) for Air, 24 h post Cl_2_ (500 ppm/20 min); 24 h post Cl2+ mitoOGG1 (n=4 each;2293 proteins).

**Supplemental Table 2**. Air vs 24 h post Cl_2_ post normalized (NSCs); n=4 (1543 proteins, both groups presented with NSCs above a zero value in ≥3 of 4 specimens for each protein analyzed, zero values were removed for statistics); there were 188 proteins that presented with statistical significance as outlined in the methods section (24 h post Cl_2_/Air; 86 increased/102 decreased).

**Supplemental Table 3.** 24 h post Cl2 vs 24 h post Cl2+mitoOGG1OGG1 study post normalized (NSCs); n=4 (1480 proteins, both groups presented with NSCs above a zero value in ≥3of4 specimens for each protein analyzed, zero values were removed for statistics), there were 118 proteins that presented with statistical significance as outlined in the methods section (24 h postCl2+mitoOGG1/24 h post Cl2; 48 increased/70 decreased).

**Supplemental Table 4.** List of primers.

## References

1. Addis DR, Aggarwal S, Doran SF, Jian MY, Ahmad I, Kojima K, Ford DA, Matalon S and Mobley JA. Vascular permeability disruption explored in the proteomes of mouse lungs and human microvascular cells following acute bromine exposure. Am J Physiol Lung Cell Mol Physiol 319: L337–L359, 2020.

2. Addis DR, Aggarwal S, Lazrak A, Jilling T and Matalon S. Halogen-Induced Chemical Injury to the Mammalian Cardiopulmonary Systems. Physiology (Bethesda ) 36: 272–291, 2021.

3. Addis DR, Lambert JA, Ford DA, Jilling T and Matalon S. Halogen gas exposure: toxic effects on the parturient. Toxicol Mech Methods 1–16, 2020.

4. Aggarwal S, Ahmad I, Lam A, Carlisle MA, Li C, Wells JM, Raju SV, Athar M, Rowe SM, Dransfield MT and Matalon S. Heme scavenging reduces pulmonary endoplasmic reticulum stress, fibrosis, and emphysema. JCI Insight 3: 2018.

5. Aggarwal S, Lazrak A, Ahmad I, Yu Z, Bryant A, Mobley JA, Ford DA and Matalon S. Reactive species generated by heme impair alveolar epithelial sodium channel function in acute respiratory distress syndrome. Redox Biol 36: 101592, 2020.

6. Agod Z, Fekete T, Budai MM, Varga A, Szabo A, Moon H, Boldogh I, Biro T, Lanyi A, Bacsi A and Pazmandi K. Regulation of type I interferon responses by mitochondria-derived reactive oxygen species in plasmacytoid dendritic cells. Redox Biol 13: 633–645, 2017.

7. Ahmad S, Matalon S and Kuebler WM. Understanding COVID-19 susceptibility and presentation based on its underlying physiology. Physiol Rev 102: 1579–1585, 2022.

8. Alishlash AS, Sapkota M, Ahmad I, Maclin K, Ahmed NA, Molyvdas A, Doran S, Albert CJ, Aggarwal S, Ford DA, Ambalavanan N, Jilling T and Matalon S. Chlorine inhalation induces acute chest syndrome in humanized sickle cell mouse model and ameliorated by postexposure hemopexin. Redox Biol 44: 102009, 2021.

9. Back SH. Roles of the Translation Initiation Factor eIF2alpha Phosphorylation in Cell Structure and Function. Cell Struct Funct 45: 65–76, 2020.

10. Banoth B and Cassel SL. Mitochondria in innate immune signaling. Transl Res 202: 52–68, 2018.

11. Bennett JA, Mastrangelo MA, Ture SK, Smith CO, Loelius SG, Berg RA, Shi X, Burke RM, Spinelli SL, Cameron SJ, Carey TE, Brookes PS, Gerszten RE, Sabater-Lleal M, de Vries PS, Huffman JE, Smith NL, Morrell CN and Lowenstein CJ. The choline transporter Slc44a2 controls platelet activation and thrombosis by regulating mitochondrial function. Nat Commun 11: 3479, 2020.

12. Bliksoen M, Mariero LH, Torp MK, Baysa A, Ytrehus K, Haugen F, Seljeflot I, Vaage J, Valen G and Stenslokken KO. Extracellular mtDNA activates NF-kappaB via toll-like receptor 9 and induces cell death in cardiomyocytes. Basic Res Cardiol 111: 42, 2016.

13. Bonetto G, Corradi M, Carraro S, Zanconato S, Alinovi R, Folesani G, Da DL, Mutti A and Baraldi E. Longitudinal monitoring of lung injury in children after acute chlorine exposure in a swimming pool. Am J Respir Crit Care Med 174: 545–549, 2006.

14. Bradley JM, Li Z, Organ CL, Polhemus DJ, Otsuka H, Islam KN, Bhushan S, Gorodnya OM, Ruchko MV, Gillespie MN, Wilson GL and Lefer DJ. A novel mtDNA repair fusion protein attenuates maladaptive remodeling and preserves cardiac function in heart failure. Am J Physiol Heart Circ Physiol 314: H311–H321, 2018.

15. Carlisle M, Lam A, Svendsen ER, Aggarwal S and Matalon S. Chlorine-induced cardiopulmonary injury. Ann N Y Acad Sci 2016.

16. Clark KA, Karmaus WJ, Mohr LC, Cai B, Balte P, Gibson JJ, Ownby D, Lawson AB, Vena JE and Svendsen ER. Lung Function before and after a Large Chlorine Gas Release in Graniteville, South Carolina. Ann Am Thorac Soc 13: 356–363, 2016.

17. Dobson AW, Grishko V, LeDoux SP, Kelley MR, Wilson GL and Gillespie MN. Enhanced mtDNA repair capacity protects pulmonary artery endothelial cells from oxidant- mediated death. Am J Physiol Lung Cell Mol Physiol 283: L205–L210, 2002.

18. Gao Y, Zhang H, Wang J, Li F, Li X, Li T, Wang C, Li L, Peng R, Liu L, Cui W, Zhang S and Zhang J. Annexin A5 ameliorates traumatic brain injury-induced neuroinflammation and neuronal ferroptosis by modulating the NF-kB/HMGB1 and Nrf2/HO-1 pathways. Int Immunopharmacol 114: 109619, 2023.

19. Gimblet GR, Houson HA, Whitt J, Reddy P, Copland JA, Kenderian SS, Szkudlinski MW, Jaskula-Sztul R and Lapi SE. PET Imaging of Differentiated Thyroid Cancer with TSHR-Targeted [(89)Zr]Zr-TR1402. Mol Pharm 21: 3889–3896, 2024.

20. Gorguner M, Aslan S, Inandi T and Cakir Z. Reactive airways dysfunction syndrome in housewives due to a bleach-hydrochloric acid mixture. Inhal Toxicol 16: 87–91, 2004.

21. Grishko V, Solomon M, Breit JF, Killilea DW, LeDoux SP, Wilson GL and Gillespie MN. Hypoxia promotes oxidative base modifications in the pulmonary artery endothelial cell VEGF gene. FASEB J 15: 1267–1269, 2001.

22. Grishko V, Solomon M, Wilson GL, LeDoux SP and Gillespie MN. Oxygen radical- induced mitochondrial DNA damage and repair in pulmonary vascular endothelial cell phenotypes. Am J Physiol Lung Cell Mol Physiol 280: L1300–L1308, 2001.

23. Harrington JS, Huh JW, Schenck EJ, Nakahira K, Siempos II and Choi AMK. Circulating Mitochondrial DNA as Predictor of Mortality in Critically Ill Patients: A Systematic Review of Clinical Studies. Chest 156: 1120–1136, 2019.

24. Hashizume M, Mouner M, Chouteau JM, Gorodnya OM, Ruchko MV, Potter BJ, Wilson GL, Gillespie MN and Parker JC. Mitochondrial-targeted DNA repair enzyme 8- oxoguanine DNA glycosylase 1 protects against ventilator-induced lung injury in intact mice. Am J Physiol Lung Cell Mol Physiol 304: L287–L297, 2013.

25. Hashizume M, Mouner M, Chouteau JM, Gorodnya OM, Ruchko MV, Wilson GL, Gillespie MN and Parker JC. Mitochondrial Targeted Endonuclease III DNA Repair Enzyme Protects against Ventilator Induced Lung Injury in Mice. Pharmaceuticals (Basel) 7: 894–912, 2014.

26. Hoyle GW and Svendsen ER. Persistent effects of chlorine inhalation on respiratory health. Ann N Y Acad Sci 2016.

27. Jani DD, Reed D, Feigley CE and Svendsen ER. Modeling an irritant gas plume for epidemiologic study. Int J Environ Health Res 26: 58–74, 2016.

28. Jurkuvenaite A, Benavides GA, Komarova S, Doran SF, Johnson M, Aggarwal S, Zhang J, Darley-Usmar VM and Matalon S. Upregulation of autophagy decreases chlorine-induced mitochondrial injury and lung inflammation. Free Radic Biol Med 85: 83–94, 2015.

29. Kuck JL, Obiako BO, Gorodnya OM, Pastukh VM, Kua J, Simmons JD and Gillespie MN. Mitochondrial DNA damage-associated molecular patterns mediate a feed-forward cycle of bacteria-induced vascular injury in perfused rat lungs. Am J Physiol Lung Cell Mol Physiol 308: L1078–L1085, 2015.

30. Kung CT, Hsiao SY, Tsai TC, Su CM, Chang WN, Huang CR, Wang HC, Lin WC, Chang HW, Lin YJ, Cheng BC, Su BY, Tsai NW and Lu CH. Plasma nuclear and mitochondrial DNA levels as predictors of outcome in severe sepsis patients in the emergency room. J Transl Med 10: 130, 2012.

31. Lazrak A, Chen L, Jurkuvenaite A, Doran SF, Liu G, Li Q, Lancaster JR, Jr. and Matalon S. Regulation of alveolar epithelial Na+ channels by ERK1/2 in chlorine-breathing mice. Am J Respir Cell Mol Biol 46: 342–354, 2012.

32. Lazrak A, Song W, Yu Z, Zhang S, Nellore A, Hoopes CW, Woodworth BA and Matalon S. Low molecular weight hyaluronan inhibits lung epithelial ion channels by activating the calcium-sensing receptor. Matrix Biol 116: 67–84, 2023.

33. Lazrak A, Song W, Zhou T, Aggarwal S, Jilling T, Garantziotis S and Matalon S. Hyaluronan and halogen-induced airway hyperresponsiveness and lung injury. Ann N Y Acad Sci 1479: 29–43, 2020.

34. Lazrak A, Yu Z, Doran S, Jian MY, Creighton J, Laube M, Garantziotis S, Prakash YS and Matalon S. Upregulation of airway smooth muscle calcium-sensing receptor by low-molecular-weight hyaluronan. Am J Physiol Lung Cell Mol Physiol 318: L459–L471, 2020.

35. Lee YL, Obiako B, Gorodnya OM, Ruchko MV, Kuck JL, Pastukh VM, Wilson GL, Simmons JD and Gillespie MN. Mitochondrial DNA Damage Initiates Acute Lung Injury and Multi-Organ System Failure Evoked in Rats by Intra-Tracheal Pseudomonas Aeruginosa. Shock 48: 54–60, 2017.

36. Ma R, Ortiz Serrano TP, Davis J, Prigge AD and Ridge KM. The cGAS-STING pathway: The role of self-DNA sensing in inflammatory lung disease. FASEB J 2020.

37. Matalon S, Yu Z, Dubey S, Ahmad I, Stephens EM, Alishlash AS, Meyers A, Cossar D, Stewart D, Acosta EP, Kojima K, Jilling T and Mobley JA. Hemopexin reverses activation of lung eIF2alpha and decreases mitochondrial injury in chlorine-exposed mice. Am J Physiol Lung Cell Mol Physiol 326: L440–L457, 2024.

38. Mills EL, Kelly B and O’Neill LAJ. Mitochondria are the powerhouses of immunity. Nat Immunol 18: 488–498, 2017.

39. Mobley JA, Molyvdas A, Kojima K, Jilling T, Li JL, Garantziotis S and Matalon S. The SARS-CoV-2 Spike S1 Protein Induces Global Proteomic Changes in ATII-Like Rat L2 Cells that are Attenuated by Hyaluronan. bioRxiv 2022.

40. Nakahira K, Kyung SY, Rogers AJ, Gazourian L, Youn S, Massaro AF, Quintana C, Osorio JC, Wang Z, Zhao Y, Lawler LA, Christie JD, Meyer NJ, Mc Causland FR, Waikar SS, Waxman AB, Chung RT, Bueno R, Rosas IO, Fredenburgh LE, Baron RM, Christiani DC, Hunninghake GM and Choi AM. Circulating mitochondrial DNA in patients in the ICU as a marker of mortality: derivation and validation. PLoS Med 10: e1001577, 2013.

41. Patterson EK, Yao LJ, Ramic N, Lewis JF, Cepinskas G, McCaig L, Veldhuizen RA and Yamashita CM. Lung-derived mediators induce cytokine production in downstream organs via an NF-kappaB-dependent mechanism. Mediators Inflamm 2013: 586895, 2013.

42. Piantadosi CA. Mitochondrial DNA, oxidants, and innate immunity. Free Radic Biol Med 152: 455–461, 2020.

43. Queern SL, Aweda TA, Massicano AVF, Clanton NA, El SR, Sader JA, Zyuzin A and Lapi SE. Production of Zr-89 using sputtered yttrium coin targets (89)Zr using sputtered yttrium coin targets. Nucl Med Biol 50: 11–16, 2017.

44. Rachek LI, Grishko VI, Musiyenko SI, Kelley MR, LeDoux SP and Wilson GL. Conditional targeting of the DNA repair enzyme hOGG1 into mitochondria. J Biol Chem 277: 44932–44937, 2002.

45. Rodriguez-Nuevo A and Zorzano A. The sensing of mitochondrial DAMPs by non-immune cells. Cell Stress 3: 195–207, 2019.

46. Scozzi D, Cano M, Ma L, Zhou D, Zhu JH, O’Halloran JA, Goss C, Rauseo AM, Liu Z, Sahu SK, Peritore V, Rocco M, Ricci A, Amodeo R, Aimati L, Ibrahim M, Hachem R, Kreisel D, Mudd PA, Kulkarni HS and Gelman AE. Circulating mitochondrial DNA is an early indicator of severe illness and mortality from COVID-19. JCI Insight 6: 2021.

47. Shanmugiah J, Zaheer J, Im C, Kang CM and Kim JS. Comparison of PET tracing and biodistribution between (64)Cu-labeled micro-and nano-polystyrene in a murine inhalation model. Part Fibre Toxicol 21: 2, 2024.

48. Simmons JD, Freno DR, Muscat CA, Obiako B, Lee YL, Pastukh VM, Brevard SB and Gillespie MN. Mitochondrial DNA damage associated molecular patterns in ventilator-associated pneumonia: Prevention and reversal by intratracheal DNase I. J Trauma Acute Care Surg 82: 120–125, 2017.

49. Simmons JD, Lee YL, Mulekar S, Kuck JL, Brevard SB, Gonzalez RP, Gillespie MN and Richards WO. Elevated levels of plasma mitochondrial DNA DAMPs are linked to clinical outcome in severely injured human subjects. Ann Surg 258: 591–596, 2013.

50. Song W, Yu Z, Doran SF, Ambalavanan N, Steele C, Garantziotis S and Matalon S. Respiratory syncytial virus infection increases chlorine-induced airway hyperresponsiveness. Am J Physiol Lung Cell Mol Physiol 309: L205–L210, 2015.

51. Summerhill EM, Hoyle GW, Jordt SE, Jugg BJ, Martin JG, Matalon S, Patterson SE, Prezant DJ, Sciuto AM, Svendsen ER, White CW and Veress LA. An Official American Thoracic Society Workshop Report: Chemical Inhalational Disasters. Biology of Lung Injury, Development of Novel Therapeutics, and Medical Preparedness. Ann Am Thorac Soc 14: 1060–1072, 2017.

52. Van Sickle D, Wenck MA, Belflower A, Drociuk D, Ferdinands J, Holguin F, Svendsen E, Bretous L, Jankelevich S, Gibson JJ, Garbe P and Moolenaar RL. Acute health effects after exposure to chlorine gas released after a train derailment. Am J Emerg Med 27: 1–7, 2009.

53. Yadav AK, Bracher A, Doran SF, Leustik M, Squadrito GL, Postlethwait EM and Matalon S. Mechanisms and modification of chlorine-induced lung injury in animals. Proc Am Thorac Soc 7: 278–283, 2010.

54. Yadav AK, Doran SF, Samal AA, Sharma R, Vedagiri K, Postlethwait EM, Squadrito GL, Fanucchi MV, Roberts LJ, Patel RP and Matalon S. Mitigation of chlorine gas lung injury in rats by postexposure administration of sodium nitrite. Am J Physiol Lung Cell Mol Physiol 300: L362–L369, 2011.

55. Zevini A, Olagnier D and Hiscott J. Crosstalk between Cytoplasmic RIG-I and STING Sensing Pathways. Trends Immunol 38: 194–205, 2017.

56. Zhang JZ, Liu Z, Liu J, Ren JX and Sun TS. Mitochondrial DNA induces inflammation and increases TLR9/NF-kappaB expression in lung tissue. Int J Mol Med 33: 817–824, 2014.

57. Zhang Q, Raoof M, Chen Y, Sumi Y, Sursal T, Junger W, Brohi K, Itagaki K and Hauser CJ. Circulating mitochondrial DAMPs cause inflammatory responses to injury. Nature 464: 104–107, 2010.

