## Supplemental Tables for "Oxidative Injury to Lung Mitochondrial DNA is a Key Contributor for the Development of Chemical Lung Injury": Supplementary Table 4 (1).docx

| **Gene name** | **Sequence (5' to 3')** |
| --- | --- |
| **mNLRP3-RT(F)** | GCTCTGACCTCTGTGCTCAA |
| **mNLRP3-RT(R)** | GTTTACAGTCCGGGTGCAGA |
| **mTLR9-RT(F)** | TGCCGACTGGGTGTATAACG |
| **mTLR9-RT(R)** | TCTCGGTCCTCCAGACACAA |
| **mOGG1 (F)** | AGC CAT CCT CGA AGA GCA AG |
| **mOGG1 (R)** | ATA CAT GGA CAT CCA CGG GC |
| **18s (F)** | GTAACCCGTTGAACCCCATT |
| **18s (R)** | CCATCCAATCGGTAGTAGCG |
| **mGAPDH (F)** | AGGTCGGTGTGAACGGATTTG |
| **mGAPDH (R)** | TGTAGACCATGTAGTTGAGGTCA |

**Supplementary Table 4:** List of primers used for Real time PCR in this study.
